# Evolution and regulation of microbial secondary metabolism

**DOI:** 10.1101/2020.09.02.280495

**Authors:** Guillem Santamaria, Chen Liao, Zhe Wang, Kyu Rhee, Francisco Pinto, Jinyuan Yan, Joao B. Xavier

## Abstract

Microbes have disproportionate impacts on the macroscopic world. This is in part due to their ability to grow to large groups and cooperatively secrete massive amounts of secondary metabolites that impact their environment. Yet, the conditions enabling secondary metabolism without compromising primary needs remain unclear. Here we investigated the biosynthesis of rhamnolipids, a secondary metabolite that *Pseudomonas aeruginosa* makes to decrease the surface tension of surrounding liquid. Using a combination of genomics, metabolomics, transcriptomics, and mathematical modeling we show that biosynthesis of rhamnolipids from glycerol varies inconsistently across the phylogenetic tree; instead, non-producer lineages are also those worse at reducing the oxidative stress of primary glycerol metabolism. The link to oxidative stress explains the inconsistent distribution across the *P. aeruginosa* tree, adding a new layer to the regulation of rhamnolipids—a microbial secondary metabolite important for fitness in natural and clinical settings.

**Significance:** The bacterium *Pseudomonas aeruginosa* is a major source of hospital-acquired infections. This pathogen’s knack for virulence relies on its ability to multiply and secrete massive amounts of substances that overwhelm microbial competitors and weaken host defenses. It remains unclear how the bacteria conciliate their need to grow and multiply—a need at the individual-level— with their ability to secrete products—a need of the population. Here we combined genomics, metabolomics and mathematical modeling to study the biosynthesis of rhamnolipids, a surfactant that *P. aeruginosa* makes to reduce the surface tension of surrounding liquids. Our study reveals a new link between oxidative stress and rhamnolipid synthesis, which helps explain how this important bacterial product has evolved and how it persists in many lineages of pathogens.

## Introduction

Microorganisms touch practically every human activity, in harmful and beneficial ways: They cause infections (1) and help us digest food (2), make bioterrorism weapons (3) and synthesize life-saving drugs (4), contribute to climate change (5) and clean our wastewater (6). The outsize impact of microbes on the macroscopic world that surrounds them comes from their ability to multiply to populations of billions that cooperate to transform chemicals in vast amounts. Many of those chemicals are not essential to microbial growth, development, and division: they are products of secondary metabolism. But despite the importance, it remains unclear why microbes invest in secondary metabolism (8, 9).

Secondary pathways divert precursors, cofactors, and energy from primary metabolism (10), a potential fitness cost to the microorganism that should be disfavored by natural selection. Secondary metabolites secreted extracellularly provide functions to the population such as cell-cell signaling (11), fighting other microbial populations (4), invading a host (12) and scavenging nutrients (13); these functions could bring a fitness advantage to the population (14). Still, a fitness benefit to the population is insufficient to explain the evolution and maintenance of a trait costly to the individual (15). Social evolution theory could provide a quantitative framework for the evolution of microbial secondary metabolism (9): If *C* is the fitness cost to the cell making the secondary metabolite and *B* is the average benefit it brings to the population, then secondary metabolism is favored when *rB>C*, where *r* is the relatedness among different genotypes across the population (16). The relatedness *r* is often low in microbial populations due to strain mixing and frequent mutations (17, 18), and this means that the cost-to-benefit ratio (*C/B*) must also be low.

Social evolution theory, then, explains why natural selection favors secondary pathways that minimize their burden on the primary metabolism (8). But social evolution does not tell how—the molecular mechanism—microbes achieved a low cost. Many secondary products require multiple enzymes for their biosynthesis (19), and the burden of those pathways on primary metabolism is likely hard to minimize. Microorganisms evolved sophisticated regulatory networks that can sense and respond to—often various—environmental stimuli (20–23). The sophisticated regulation of secondary metabolic pathways explains conditional phenotypes in laboratory culture, such as when secondary metabolites depend on the type and quantity of the nutrients provided, especially the carbon source (24), and when production occurs in specific growth phases, typically the stationary phase (25). The molecular mechanism regulating secondary metabolism provides a level of understanding of the same biological phenomenon complementary to evolutionary theory (26).

Here we investigate both the evolution and the regulation of rhamnolipid synthesis in *Pseudomonas aeruginosa*. Rhamnolipids are secondary metabolites that decrease the surface tension of the surrounding liquid (27). A *P. aeruginosa* cell can produce 20% or more of its dry weight in rhamnolipids (28), which is potentially a huge cost to the individual cell (29). But the rhamnolipids secreted collectively by many bacteria enable spectacular social behaviors like swarming (30), climbing over walls (31), killing microbial competitors (32) and breaching the epithelial barriers of a host (33).

The first step in rhamnolipid biosynthesis is catalyzed by the RhlA enzyme, which takes a metabolic intermediate from fatty acid synthesis, β-hydroxyacyl-ACP, and produces 3-(3-hydroxyalkanoyloxy) alkanoic acids (HAAs) (34). The RhlB and RhlC enzymes each add one unit of rhamnose to the HAAs to produce mono-rhamnolipids and di-rhamnolipids (35). The expression of the *rhlAB* operon is regulated by a sophisticated network that includes at least three quorum sensing signals (36, 37) and is conditional to nutrients in the media, requiring a high carbon-to-nitrogen or carbon-to-iron ratio (29, 38, 39). This regulation produces a strategy called metabolic prudence that delays the expression of rhamnolipid biosynthetic genes to times when the population is large enough, but the individual cell also has sufficient carbon source (29). Compared to non-regulated constitutive production, the regulation of rhamnolipid biosynthesis genes make it robust against cheating by lowering the *C/B* ratio even when the relatedness *r* is relatively low (40).

Our understanding of rhamnolipids biosynthesis comes mostly from laboratory strains, and this leaves clinically relevant questions unanswered. The three genes encoding the secondary pathway—*rhlA*, *rhlB* and *rhlC*—are conserved across the *Pseudomonas* and the *Burkholderia* genera (42). However, the ability to make rhamnolipids can vary widely, even among closely related strains of *P. aeruginosa* (41). What explains this phenotypic variation?

Here we investigate the phenotypic diversity of set of *P. aeruginosa* strains isolated from patients with cancer. We use glycerol as carbon source, a relevant nutrient for *P. aeruginosa* infection (43, 44) that puts a substantial strain on *Pseudomonas* primary metabolism (45). We show that the lineages that lost their ability to make rhamnolipids from glycerol grow not less but slower on glycerol, compared to lineages capable of rhamnolipid production. Then we use metabolomics and mathematical modeling to link the inability to reduce oxidative stress produced by primary metabolism in glycerol with the lack of rhamnolipid synthesis. This link adds a new layer to the regulation of rhamnolipid synthesis and explains the inconsistent variation of the surfactant production phenotype across the phylogenetic tree.

## Results

### Surfactant activity lost in strains with impaired primary metabolism

We investigated the phenotypic variability among clinical isolates in a diverse panel of 31 *P. aeruginosa* strains, including 28 isolates from infected patients (46) and the three type strains PAO1, PA14 and PA7, all of which have their genomes sequenced (**Fig. 1A**). We grew each strain in the same synthetic media using glycerol as the sole carbon source and ammonium sulfate as the sole nitrogen source, producing a carbon-to-nitrogen molar ratio of 7.0, which enabled rhamnolipid synthesis in strain PA14 (38). We classified each strain as non- (and very low, n=8), mild- (n=6) and strong-producers (n=17) based on drop collapse assay, which assess the surfactant activity of each strain’s secretions. The strong producers showed similar surface tension to our laboratory strain PA14, non-producers showed surfactant activity indistinguishable to cell-free media, and mild-producers show activities in between. There was no obvious relation between the body site infected in the patient and the surfactant activity phenotype. There was also no coherent distribution with the isolates’ phylogeny, which we built using the sequence variation in core genes (**Fig. 1B**; also supported by a poor correlation in Moran’s I test, *P*=0.14). Ancestral reconstruction the surfactant activity trait supports that the ability to synthesize surfactants from glycerol was present in the common ancestor of all strains (**Supplementary Fig. 1**). The genes *rhlA*, *rhlB* and *rhlC*, the genes encoding the rhamnolipid biosynthesis pathway, were present in all 31 isolates, and their sequence was 100% conserved. This indicates that the absence of surfactant activity was not due to a loss of the rhamnolipid biosynthetic pathway. Instead, we hypothesized that the phenotypic variation could be caused by some lineages losing their ability by rewiring their regulatory network to overcome specific selective pressures faced during their evolution.

**Figure 1.**
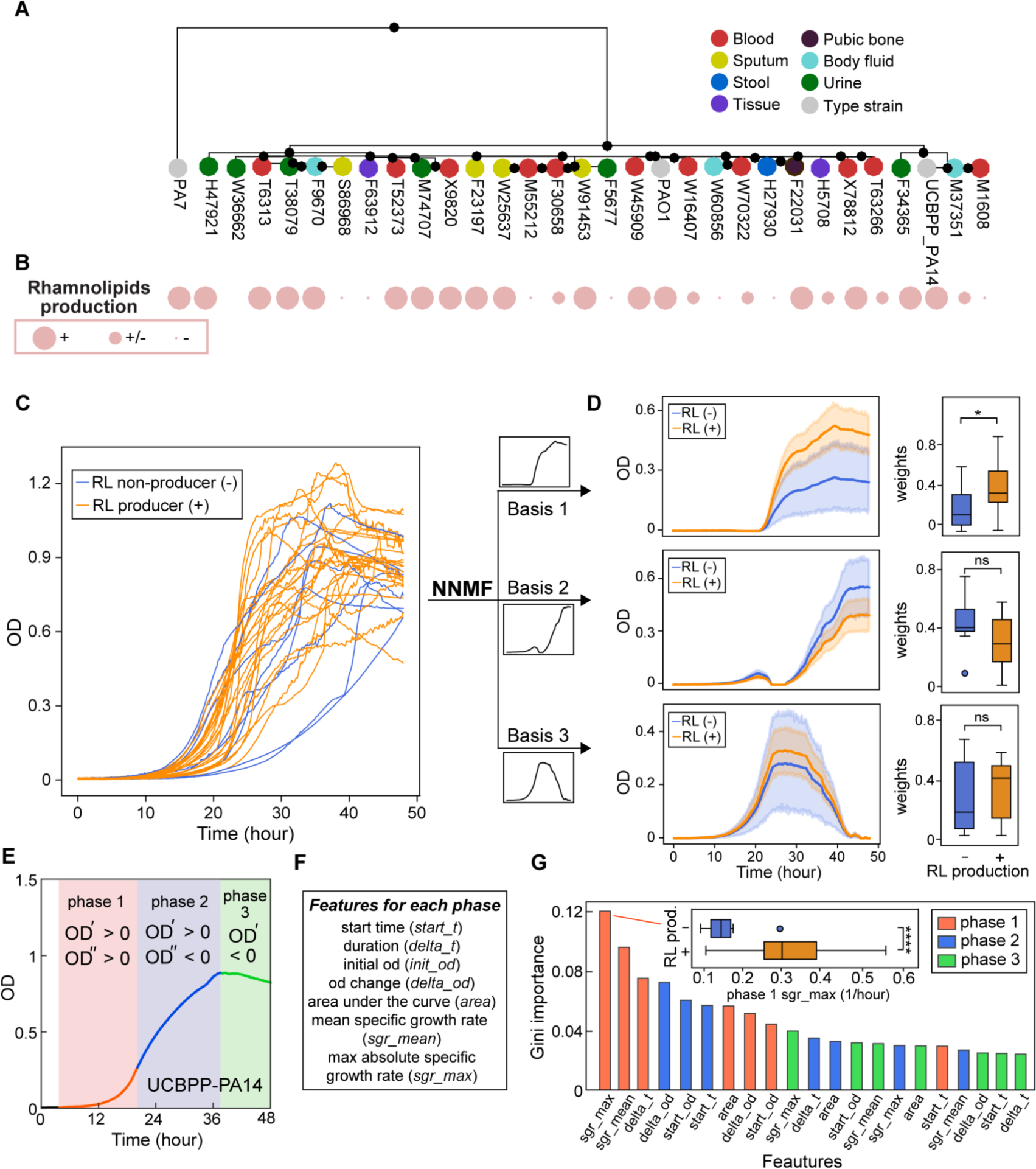
Diversity of rhamnolipid production across the *P. aeruginosa* phylogeny of core genomes and growth curve features distinguish rhamnolipid producers from non-producers. **A.** Phylogeny of clinical isolates obtained from patients with cancer at MSKCC (21, 75) together with reference type strains PAO1, PA14 and PA7. The tissue where each isolate was originally isolated is labeled by circle colors. **B**. Rhamnolipid production phenotypes. The ability of producing rhamnolipids of these strains is qualitatively indicated by circle sizes (right column). **C,D.** Unsupervised feature selection using non-negative matrix factorization (NNMF), which decomposes growth curves of all *Pseudomonas* isolates into three additive basis functions (features) such that each growth curve can be approximately represented by the weighted sum of these functions. **C**. Growth curves from both rhamnolipid producers (orange) and non-producers (blue). **D**. Decomposed components (basis function multiplied by weights; left panels) and weights (right panels) from NNMF grouped by rhamnolipid (RL) production. The shaded areas represent 95% bootstrap confidence intervals of the mean. **E-G**. Supervised feature selection using Random Forest classifier. **E,F**. Feature extraction method. Each growth curve (excluding the initial lag phase) was divided into three phases (**E**) and each phase was described by 7 quantitative features (**F**). **G.** Ranking of feature importance in classifying rhamnolipid producers. Inset: boxplot of maximum specific growth rate of phase I grouped by rhamnolipid production. Welch’s t-test was used in (**D**) and (**G**) for significance testing. ****, p-value ≤ 0.0001; *, p-value ≤0.05; ns, p-value > 0.05.

**Figure 2.**
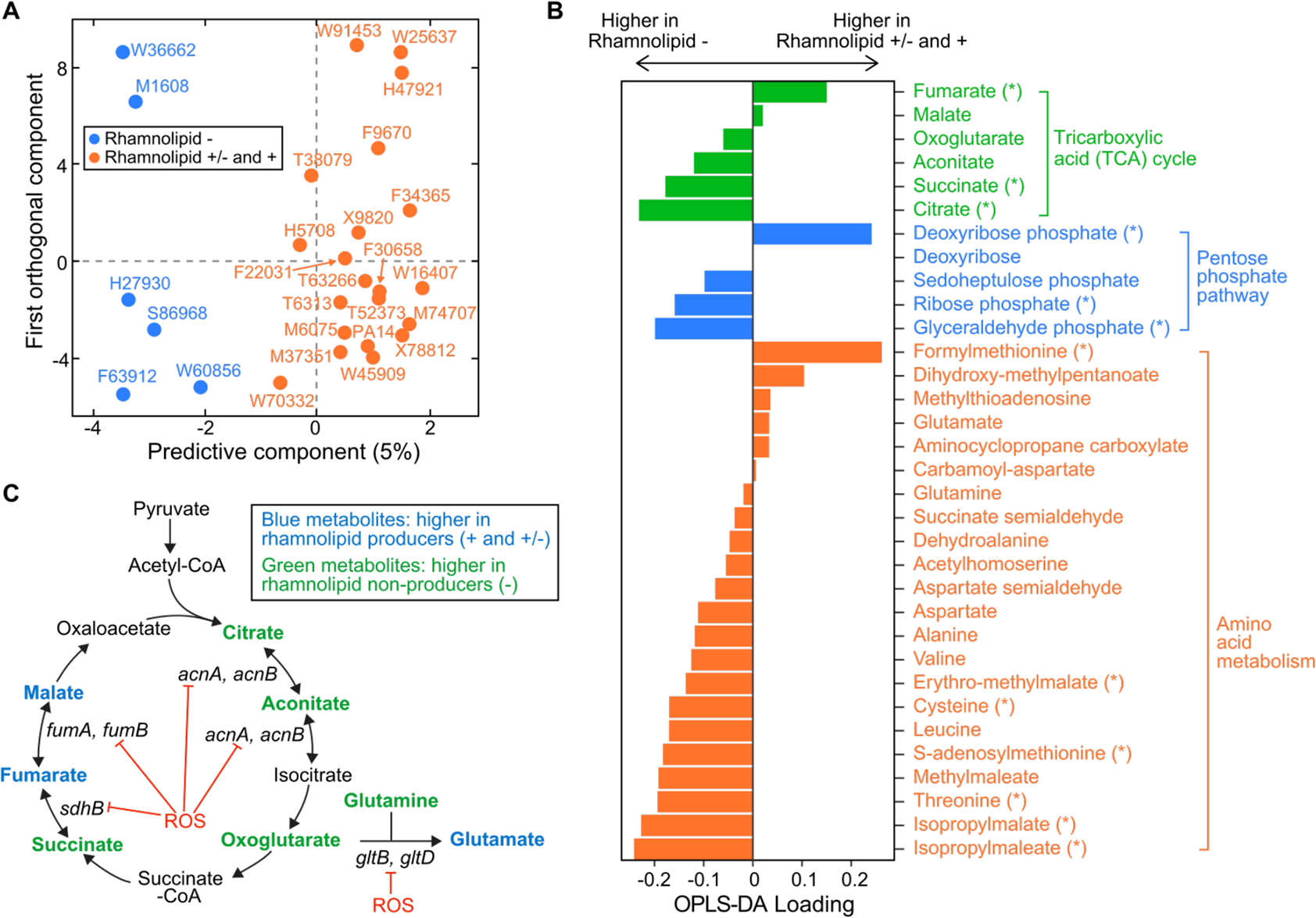
Comparative metabolomics between rhamnolipid producers (include both mild and strong production levels) and non-producers. **A.** The scores plot of the OPLS-DA (orthogonal partial least squares discriminant analysis) model. The first principle component separated the rhamnolipid non-producers from producers. **B.** OPLS-DA loading values for the first principle component of a selected number of metabolites. Metabolites with asterisk (*) are significantly different between producers and non-producers (adjusted p*-*value of a Mann-Whitney test < 0.05). **C**. Differential abundance of metabolites involved in reactions catalyzed by some Fe-S-containing enzymes whose activities are inhibited by reactive oxygen species (ROS). Abbreviations: *acnA* (aconitate hydratase A), *acnB* (aconitate hydratase B), *sdhB* (succinate dehydrogenase subunit), *fumA* (fumarase A), *fumB* (fumarase B), *gltB* (glutamate synthase subunit), *gltD* (glutamate synthase subunit).

We investigated whether variation in the accessory genome (the set of genes missing in at least one strain) could explain the phenotype differences. Among the 8,290 accessory genes, 354 are only absent in those non-producers but present in all mild- and strong-producers (**Supplementary Table 1**). A principal component analysis of the presence-absence profiles showed that our isolates are not grouped by their surfactant phenotype (**Supplementary Fig. 2**), consistent with the conclusion drawn from the core genome.

To search for a general explanation, we conducted analyses that showed producers have faster primary metabolisms in the same media of drop collapse assay. This finding was unexpected because, if anything, secondary metabolism should divert resources from primary metabolism (10): strains that produce surfactants would grow slower, not faster. We started by noting intriguing differences in the growth curves of the 31 strains: although there were no notable differences in the population densities reached, the cultures varied widely in their lag-phase and the exponential phase (**Fig. 1C**). The shape of the growth curve, as a whole, was not predictive of surfactant activity: hierarchical clustering using the entire time series failed to separate producers and non-producers (**Supplementary Fig. 3**). But local differences revealed that it’s how fast a strain grows, not how much it grows, that distinguishes producers from non-producers. Non-negative matrix factorization (47) decomposed each growth curve as a weighted sum of three basis functions (i.e., features) and showed that—although the growth curves of the rhamnolipid producers (orange lines; including mild- and strong-producers) and non-producers (blue lines) largely overlapped (**Fig. 1C**)—producers had higher weights for base 1 which agrees with a faster growth rate in the exponential phase (**Fig. 1D**). This was also confirmed by dividing each growth curve into three phases (**Fig. 1E** and **Supplementary Fig. 4**) and defining 7 quantitative features to characterize each growth phase (**Fig. 1F** and **Supplementary Table 2**). Random Forest classification revealed that the top two features associated with rhamnolipid production were the maximum and the average specific growth rates in phase I, the exponential growth phase (**Fig. 1G**). Features of phase III (the decay phase) such as the phase starting OD which represents the final biomass produced by different strains in phase II, were among the least significant.

These analyses suggested that the biosynthesis of rhamnolipids, which peaks when growth slows down from exponential (37–39), is associated with the speed of the exponential phases and hence an efficient primary metabolism on glycerol. The result supports the idea that the non-producer lineages had particular life histories that selected for specific metabolic adaptations, and that those adaptations had the collateral damage of impacting their ability to grow on glycerol as a sole carbon source.

### Metabolomics shows differences in intracellular metabolomes associated with loss of surfactant biosynthesis

We then investigated metabolomic differences that distinguish producer and non-producer lineages. We extracted the intracellular metabolome of all our strains except for two non-producers (M55212 and F23197), which grew too slowly during the transition between phase I and phase II when rhamnolipid production begins (**Materials and Methods**). LC-MS (Liquid chromatography-Mass spectrometry) revealed 67 metabolites whose identities are known, and their abundances varied significantly across the tested strains (**Supplementary Fig. 5A-B**). Hierarchical clustering and principal component analysis (**Supplementary Fig. 6A-B**) showed a reasonable separation between producers and non-producers. Fitting the data using the Orthogonal Projections to Latent Structures-Discriminant Analysis (OPLS-DA) (48) confirmed a significant correlation between metabolite amount and the surfactant phenotype (**Fig. 3A**, R^2^ = 0.82, Q^2^ = 0.66, *p-*value = 5e-4). A Mann-Whitney U test revealed 15 such metabolites statistically significantly different in producers vs non-producers, which we then used to identify pathways perturbed in non-producers using the FELLA algorithm for pathway enrichment analysis (49) (**Supplementary Fig. 7**, **Supplementary Table 3**). The most perturbed pathways were the TCA cycle and amino acid metabolism (**Fig. 3B**). Three out of six metabolites in the TCA cycle—fumarate, succinate and citrate—were significantly changed: fumarate and malate had positive loadings, while those of citrate, succinate, and to a lesser extent cis-aconitate and alpha-ketoglutarate were negative (**Fig. 3B**). Pyruvate remained relatively constant across all the strains (**Supplementary Fig. 5A**), implying that the differential responses in the TCA cycle were independent from the changes in its upstream central carbon metabolism.

**Figure 3.**
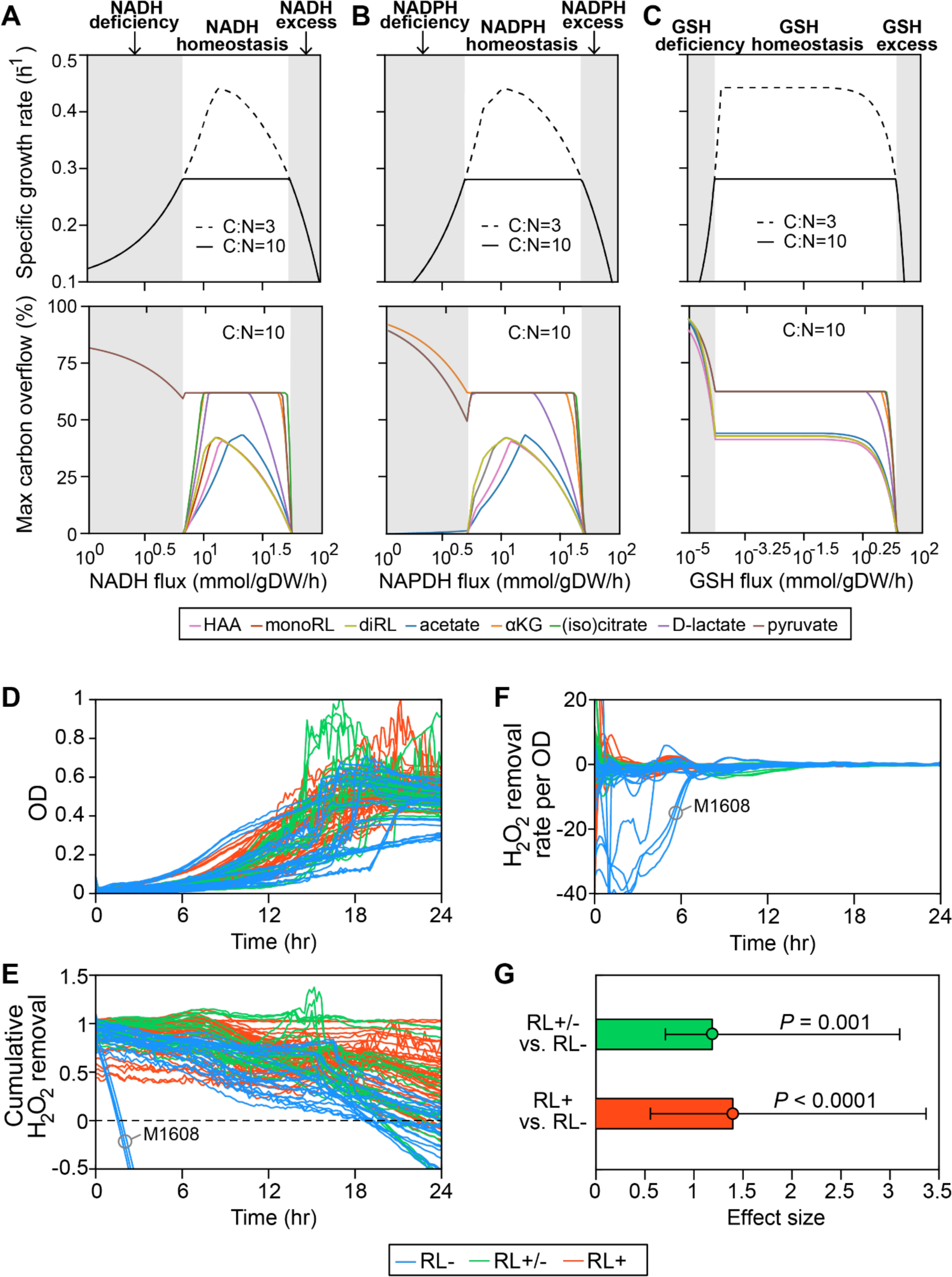
Dependence of rhamnolipid secretion on intracellular redox status. **A-C**. Model simulation. The redox stress levels are perturbed by altering fluxes of (**A)** NADH (reduced nicotinamide adenine dinucleotide), (**B**) NADPH (reduced nicotinamide adenine dinucleotide phosphate) and (**C**) GSH (reduced glutathione). Upper panels are predicted maximum growth rates and lower panels are predicted maximum byproduct secretion fluxes. C:N indicates the carbon-to-nitrogen influx ratio provided by culture medium. C:N=3 and C:N=10 represent carbon- and nitrogen-limiting conditions respectively. Abbreviations: HAA: 3-(3-hydroxyalkanoyloxy) alkanoate; monoRL: monorhamnolipid; diRL: dirhamnolipid; aKG: alpha-ketoglutarate. **D-G**. Experimental validation via comparison of the ability of removing culture medium hydrogen peroxide (H_2_O_2_) among strong rhamnolipid producers (+), weak producers (+/-), and non-producers (-). **D.** Population density. **E.** The total amount of hydrogen peroxide removed from the environment. Negative values indicate net cellular production of hydrogen peroxide released to the environment. **F.** The specific hydrogen peroxide removal rate. In both **E** and **F**, each trajectory of H_2_O_2_ fluorescence intensity was normalized to the averaged trajectory of the wild-type UCBPP-PA14 strain. **G.** Effect size of rhamnolipid production as a predictor of H_2_O_2_ removal rate per OD in a mixed-effect analysis.

Besides the TCA cycle metabolites, most of the annotated compounds in the metabolism of branched chain amino acids (leucine/isoleucine, valine) and sulfur-containing amino acids (cysteine/methionine) were more abundant in non-producers than producers (**Supplementary Fig. 7**). A notable exception was formylmethionine (fMet), which was lower in non-producers. We went back to previous metabolomics data (50) and found that the Δ*rhlA* mutant of *P. aeruginosa* PA14 also had lower levels of fMet compared with its wild-type (**Supplementary Fig. 8**). Since Δ*rhlA* mutant grows just as fast in glycerol as the wild-type (51) the link between lower fMet and lack of rhamnolipid production should not be simply due to a growth defect.

### Theory formalizes the link between oxidative stress, slower growth and loss of rhamnolipid synthesis

Several theoretical arguments support that non-producers might have slower primary metabolisms because they suffer from higher oxidative stress than the producers. First, the major differences include TCA intermediates (**Fig. 3B**). The TCA cycle harbors five enzymes with Fe-S clusters, aconitase A, aconitase B, succinate dehydrogenase, fumarase A, fumarase B (52), which are among the most vulnerable to oxidative stress. If non-producers are less successful with removing the oxidative stress from growth in glycerol, then the stress accumulated could reduce flux through the TCA cycle, which would slow down growth. The accumulation of succinate and depletion of fumarate in non-producers could be due to a reduced activity of succinate dehydrogenase (SDH) under oxidative stress: SDH is a membrane-bound dehydrogenase linked to the respiratory chain—a major site of ROS production in the cell—and also a member of the TCA cycle that catalyzes the oxidation of succinate into fumarate (53).

Since SDH contains [2Fe-2S], [3Fe-4S], and [4Fe-4S] clusters (54), ROS that damages Fe-S clusters could decrease SDH activity *in vivo*.

Second, we saw that non-producers also included intermediate metabolites in amino acid biosynthetic pathways which are also substrates of Fe-S containing enzymes (**Fig. 3B**). For example, the glutamate synthetase has a large chain (encoded by *gltB*) and a small chain (encoded by *gltD*), and both subunits contain Fe-S clusters that catalyze the production of glutamate from oxoglutarate and glutamine (55). Non-producers had slightly higher oxoglutarate and glutamine as well as lower glutamate than producers.

Finally, we used a whole-genome reconstruction to model the intracellular metabolic fluxes during growth in the glycerol medium (56, 57) which supported our observations. Rhamnolipids are produced when carbon is in excess (38, 39). In the simulations, rhamnolipids secretion occurs when C:N flux ratio > 6.3 (**Supplementary Fig. 9**). We then set C:N to 10.0, which exceeds the minimum threshold, and we simulated various levels of redox stress level by changing the flux levels of three redox molecules—NADH (reduced nicotinamide adenine dinucleotide), NADPH (reduced nicotinamide adenine dinucleotide phosphate) and GSH (reduced glutathione)—responsible for the bulk of cellular electron transfer and the main sources of reactive oxygen species (ROS) (58). Although redox stress is ultimately caused by high ROS levels, not fluxes, and ROS was not an explicit variable in our model, we assumed that ROS can perturb intracellular metabolic fluxes, particularly the fluxes of redox molecules, and we then asked how cell growth and rhamnolipid secretion responded to these ROS-mediated indirect perturbations.

For all three redox molecules, we found that the maximum growth rate was maintained at an intermediate flux range (redox homeostasis; **Fig. 4A-C**, upper panels, white area). The growth rate decreased gradually as the flux of redox molecules changed in either direction away from this homeostatic interval (gray shading). The reason for the gradual decrease in growth rate is that deficiency or excess of these redox molecules which participate in the redox control of a great variety of biological processes disturbs the primary metabolism for growth. Importantly, before the growth starts slowing down due to an excess in redox species there was an abrupt shutdown in the overflow on carbon into many central carbon metabolites and—importantly— in the secretion of HAAs, mono-and di-rhamnolipids (**Fig. 4A-C**, lower panels). The shutdown of overflow reactions under excess of redox molecules mimics the metabolic state in the rhamnolipid non-producers who lose metabolic plasticity to overflow metabolites without compromising growth. When we forced these secretion fluxes to zero—to simulate a Δ*rhlA* mutant—the simulation did not predict a growth benefit. This result agrees with data showing that the *ΔrhlA* mutant of *P. aeruginosa* strain PA14 has no growth benefit in glycerol media compared with wild-type (51). This also suggests that regaining rhamnolipid secretion should not restore fast growth in glycerol to non-producers, which agrees with the idea that loss of rhamnolipid synthesis is a consequence—not a cause—of the slower growth in non-producers. Notably, most of the overflowed metabolites are carbon-rich molecules, including HAA, mono- and di-rhamnolipid (59). Collectively, these theoretical arguments support a link between rhamnolipids secretion and oxidative stress, where fast growth and rhamnolipid secretion are both impacted by the accumulation of ROS produced by primary metabolism.

**Figure 4.**
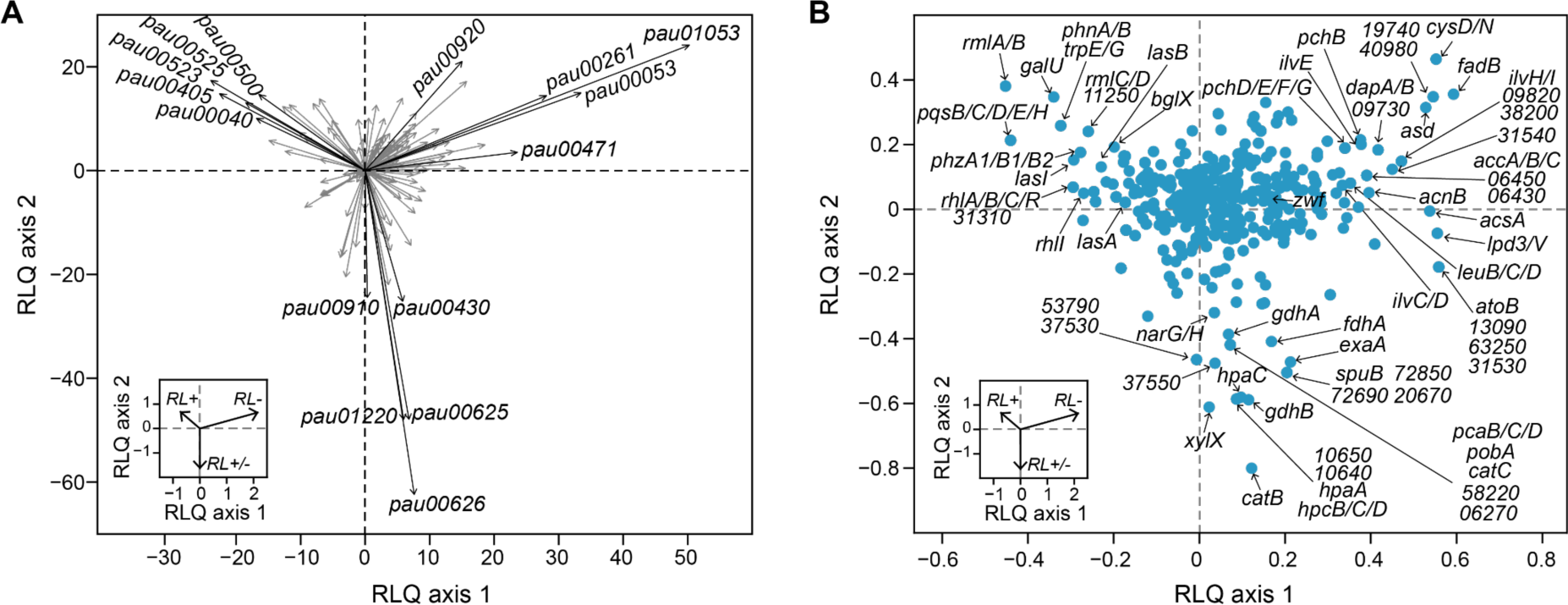
Functional pathways (**A**) and individual gene expressions (**B**) associated with rhamnolipid production. RLQ axes 1 and 2 are the first two axes of RLQ analysis. The insets in both A and B represent the rhamnolipid synthesis phenotypes in the principal RLQ axes. Genes and pathways aligning with the direction of each phenotypic category are associated with the phenotype. In panel A, the top 5 pathways (“pau” are KEGG pathway IDs) associated with each phenotype are highlighted in black. The five-digit number in panel B represents the PA14 locus ID. RL+: strong rhamnolipid producer; RL+/-: mild-rhamnolipid producer; RL-: rhamnolipid non-producer. Description of the KEGG pathways: pau01053 (Biosynthesis of siderophore group nonribosomal peptides), pau00053 (Ascorbate and aldarate metabolism), pau00261 (Monobactam biosynthesis), pau00471 (D-Glutamine and D-glutamate metabolism), pau00920 (Sulfur metabolism), pau00626 (Naphthalene degradation), pau01220 (Degradation of aromatic compounds), pau00625 (Chloroalkane and chloroalkene degradation), pau00430 (Taurine and hypotaurine metabolism), pau00910 (Nitrogen metabolism), pau00525 (Acarbose and validamycin biosynthesis), pau00405 (Phenazine biosynthesis), pau00523 (Polyketide sugar unit biosynthesis), pau00500 (Starch and sucrose metabolism), pau00040 (Pentose and glucuronate interconversions).

**Figure 5.**
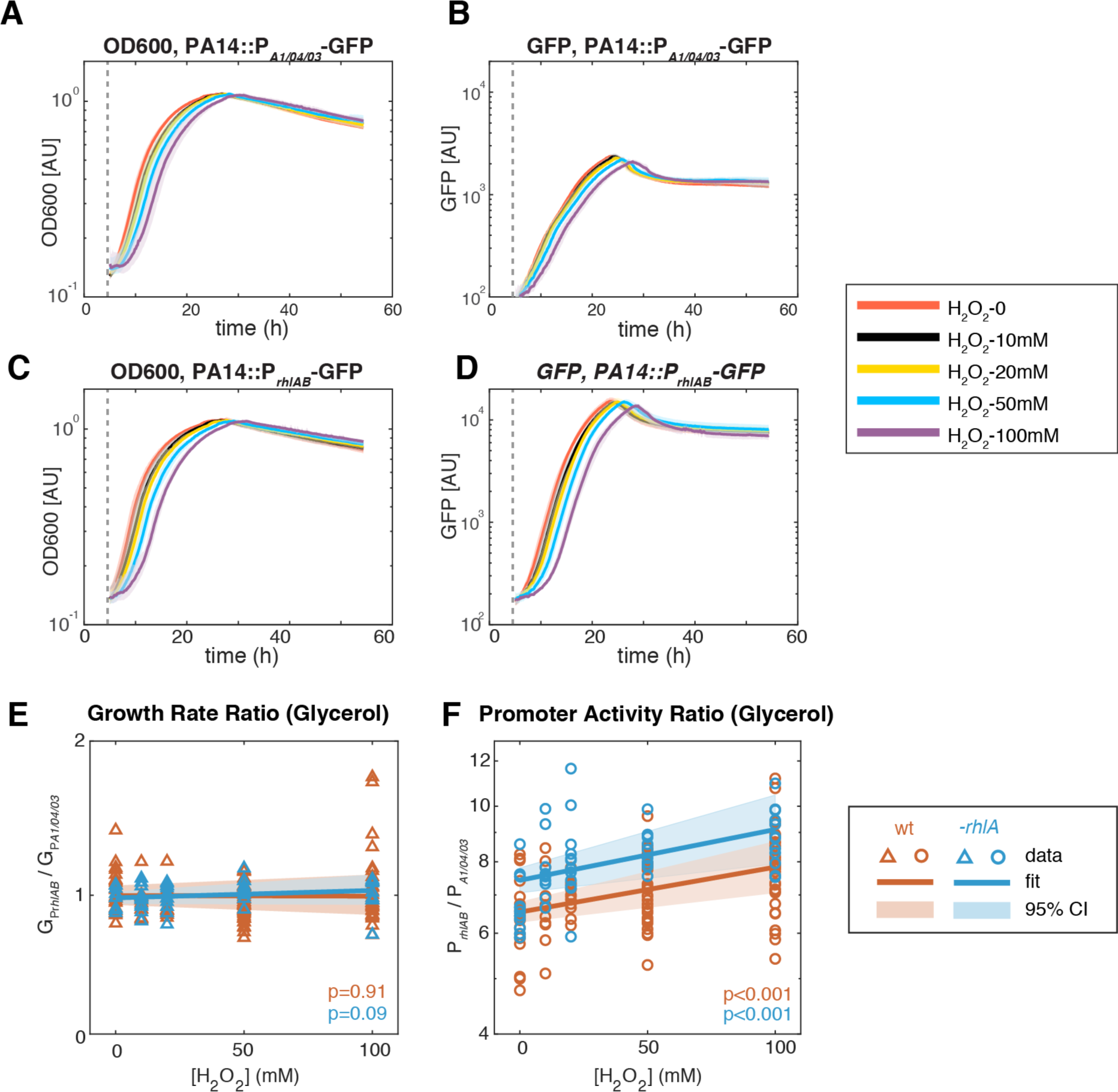
Comparing the effect of different oxidative stress on the *rhlAB* gene expression and cell growth in synthetic media of different carbon sources. (A-D) Demonstration of the cell growth (A, C) and GFP expression (B, D) of PA14::P_*A1/04/03*_-GFP (constitutive promoter) and PA14::P_*rhlAB*_-GFP when different levels of H_2_O_2_ was added to the media at the beginning of exponential growth (gray dash line) using one experiment. The shaded area represents the range of all replicates from the same experiment. (E, F) The ratio of growth rate (E) and promoter activity (F) of PA14::P_*rhlAB*_-GFP to PA14::P_*A1/04/03*_-GFP in phase II under different levels of H_2_O_2_ when glycerol was used as the carbon source of the synthetic media. The p-value represents how significant the coefficient (slope) is using fitglme. Each dot represents one replicate.

### Experiments confirm that non-producers are worse at reducing oxidative stress

Our model suggested that non-producers suffer from higher oxidative stress. *Pseudomonas* have a variety of antioxidant enzymes including catalases, glutathione reductase, NADH peroxidase and cysteine-based peroxidases (60), which were only secreted when the growth rate maintained its maximum value under excess carbon, a finding also consistent with experiments in PA14 and PAO1 (29, 38, 39). Isogenic mutants lacking some of those mechanisms are impaired in rhamnolipid synthesis (61). If the slower growth in non-producers is due to impairments in some of those mechanisms this could affect surfactant biosynthesis. To test this hypothesis we compared the dynamics of hydrogen peroxide (H_2_O_2_) (**Fig. 3D,E**). H_2_O_2_ is a representative ROS that can diffuse freely between cells and the environment; we can access its dynamics by measuring the fluorescence by the Amplex assay, where the difference between cell cultures and baseline reveals each strain’s net ability to reduce H_2_O_2_ (62). All strains except for M1608 quenched the signal over the first 18 hours (**Fig. 3E**). As expected, the non-producers (blue lines) removed less H_2_O_2_ than mild rhamnolipid producers (green lines) and strong producers (red lines). To factor out the possibility that the non-producers performed worse due to lower cell density, we calculated the H_2_O_2_ removal rate per cell and observed a similar relative trend of H_2_O_2_ degradation between non-producers and producers (**Fig. 3F**): a linear mixed-effect model quantified the difference that mild and strong producers have in per-cell H_2_O_2_ removal rates, which significantly higher than in non-producers (*P* < 0.001; **Fig. 3G**). This supports that rhamnolipid non-producers are suffering from primary metabolism generated oxidative species, which can be caused by high production or slow degradation. Interestingly, the worst strain in this assay, M1608, lacks *katE* (**Supplementary Table 1**), a catalase that degrades H_2_O_2_ (63). Another non-producer, F5677, lacks the redox sensor *soxR (64)*.

### RNAseq reveals oxidative stress response in non-producers

To understand the gene expression changes in rhamnolipid non-producers compared to the producers, we selected 11 representative strains (6 strong producers, 3 mild producers and 2 non-producers), in proportion to the number in each category. We then assess their transcriptomics in Phase II when rhamnolipids biosynthesis starts in the producers (**Materials and Methods**). A PCA of the gene expression profiles could not cluster the strains according to their surfactant phenotypes (**Supplementary Fig. 10**). We then turned to RLQ analysis (**Materials and Methods**) to find both genes and pathways associated with rhamnolipid production. At the gene function level, the top 5 pathways enriched in each group do not overlap: The strong producers were enriched in acarbose and validamycin biosynthesis (pau00525), phenazine biosynthesis (pau00405), polyketide sugar unit biosynthesis (pau00523), starch and sucrose metabolism (pau00500), and pentose and glucuronate interconversions (pau00040) (**Fig. 4A**, **Supplementary Table 4**). By contrast, biosynthesis of siderophore group nonribosomal peptides (pau01053), ascorbate and aldarate metabolism (pau00053), monobactam biosynthesis (pau00261), D-glutamine and D-glutamate metabolism (pau00471), and sulfur metabolism (pau00920) were the top 5 pathways enriched in the non-producers (**Fig. 4A**, **Supplementary Table 4**).

As expected, the genes in the biosynthesis of rhamnose (*rmlA, rmlB, rmlC, rmlD*), rhamnolipid biosynthesis (*rhlA, rhlB, rhlC, rhlR, rhlI*), quorum sensing (*lasA, lasB, lasI, pqsABCDEH*) were significantly lower in the non-producers (**Fig. 4B**, **Supplementary Fig. 11**). Interestingly, *in vitro* data shows that the expression of genes from the above list, including *lasI, lasR, rhlI, rhlR, pqsA, pqsR*, in *P. aeruginosa* PAO1, can be significantly reduced by quorum sensing quencher and ROS molecule H_2_O_2_ (65), supporting that the loss of the phenotype may be linked to oxidative stress. More importantly, the *zwf* gene encoding glucose 6-phosphate dehydrogenase (G6PDH) was expressed higher in non-producers (**Supplementary Fig. 11**). G6PDH is a major source of NADPH and contributes to antioxidant defenses by reducing oxidized glutathione, and its higher expression supports our hypothesis that non-producers experience higher oxidative stress. Other genes including the peroxide-sensing transcriptional regulator (*oxyR*), catalases (*katA, katB*), alkyl hydroperoxide reductases (*ahpC, ahpF*), glutaredoxin (*grx*), glutathione reductase (*gor*), and thioredoxins (*trxA, trxB1, trxB2*) were not highly expressed in the producers (**Supplementary Fig. 11**), suggesting that the antioxidant defense may be regulated at the metabolic level.

### Exogenous oxidative stress induces rhamnolipid biosynthesis in a strain that can deal with stress produced by primary metabolism

So far, we have shown that surfactant producers reduce oxidative species better, and we provided a model to explain the link between a faster exponential growth and the ability to make rhamnolipids, indicating that rhamnolipid production may be co-evolving with primary metabolism through the regulation of oxidative stress. Another proof of this co-evolving mechanism is that rhamnolipid production evolves to be part of a wide oxidative stress response. This possibility was suggested by experiments showing that the induction of rhamnolipids synthesis in an engineered strain of *P. putida* reduced the release of CO_2_, which was produced using O_2_ as electron acceptor, suggesting that rhamnolipids could function as a secondary electron sink (66). To investigate this possibility, we tracked gene expression in one of the rhamnolipid producers—the strain PA14—using a reporter fusion (*PrhlAB*-GFP) to track the expression of *rhlAB* after adding different levels of H_2_O_2_ (**Fig. 5**, **Supplementary Fig. 12**). We added H_2_O_2_ when the cells started entering the exponential phase, instead of in the middle of the exponential phase, to avoid strong perturbation of the measuring signal. Indeed, H_2_O_2_ delayed cell growth (**Figure 5A, C**) but no difference was observed for the exponential growth rate, indicating that this strain can sustain its primary metabolism despite the oxidative stress. However, the *rhlAB* promoter activity increased as H_2_O_2_ was added compared to a strain carrying the constitutive promoter (PA1/04/03) (**Fig 5F**, orange). This indicated that adding an exogenous redox stress to a strain that can deal with it—since it did not slow down the speed of cell growth in phase I and phase II—can give feedback to increase *rhlAB* expression in the PA14 strain. We next compared this with the strain *ΔrhlA* (**Supplementary Fig. 13**). We observed a similar increase of P*rhlAB-*GFP expression with increasing H_2_O_2_ in *ΔrhlA* strain (**Figure 5E,F**, blue). This suggests an attempt to increase rhamnolipid biosynthesis, which this strain is incapable of because it lacks the *rhlA* gene. However, we saw no decrease in the growth rate, indicating that the lack of rhamnolipid synthesis had no detrimental impact on primary metabolism. Interestingly, however, the baseline of the promoter activity increased significantly (1.14x, p<0.01) compared to the wild type. This higher expression is not due to the genetic background, because the promoter activity between PA14 and *ΔrhlA* was not significantly different in the absence of H_2_O_2_ (p=0.3). Taken together, these results indicate that rhamnolipid biosynthesis can be induced by additional redox stress, and that the signal for the induction is stronger when rhamnolipid synthesis fails to start, while primary metabolism is unaffected in these conditions.

## Discussion

Our results suggest an expanded model for the regulation of the rhamnolipid secondary pathway to explain the variation of surfactant activity observed in *P. aeruginosa* isolates (**Fig. 6**). A microbe can produce secondary metabolites such as rhamnolipids only after it meets its primary requirements of energy production, biomass synthesis and reduction of oxidative stress (**Fig. 6A**) and has excess carbon source. The lineages that retained their ability to synthesize rhamnolipids were also those capable of reducing the oxidative stress created by primary metabolism in glycerol, and they could overflow any excess carbon to secondary rhamnolipid metabolism (**Fig. 6B**). The loss of rhamnolipids in non-producer lineages was distributed inconsistently across the phylogenetic tree, possibly due to the life histories of each lineage. Consistent with the model, those non-producers had impaired abilities to reduce oxidative stress, a slower exponential growth indicating a deficient primary metabolism, perturbed levels of TCA cycle metabolites consistent with oxidative damage and accumulated intermediates of amino acid synthesis—also consistent with a slower primary metabolism. Those burdens prevent the cell from producing rhamnolipids as the resources can be prioritized for primary metabolism, as we observed that the OD at the end of exponential growth is not a good distinguisher for producers and non-producers (**Fig. 6C**).

**Figure 6.**
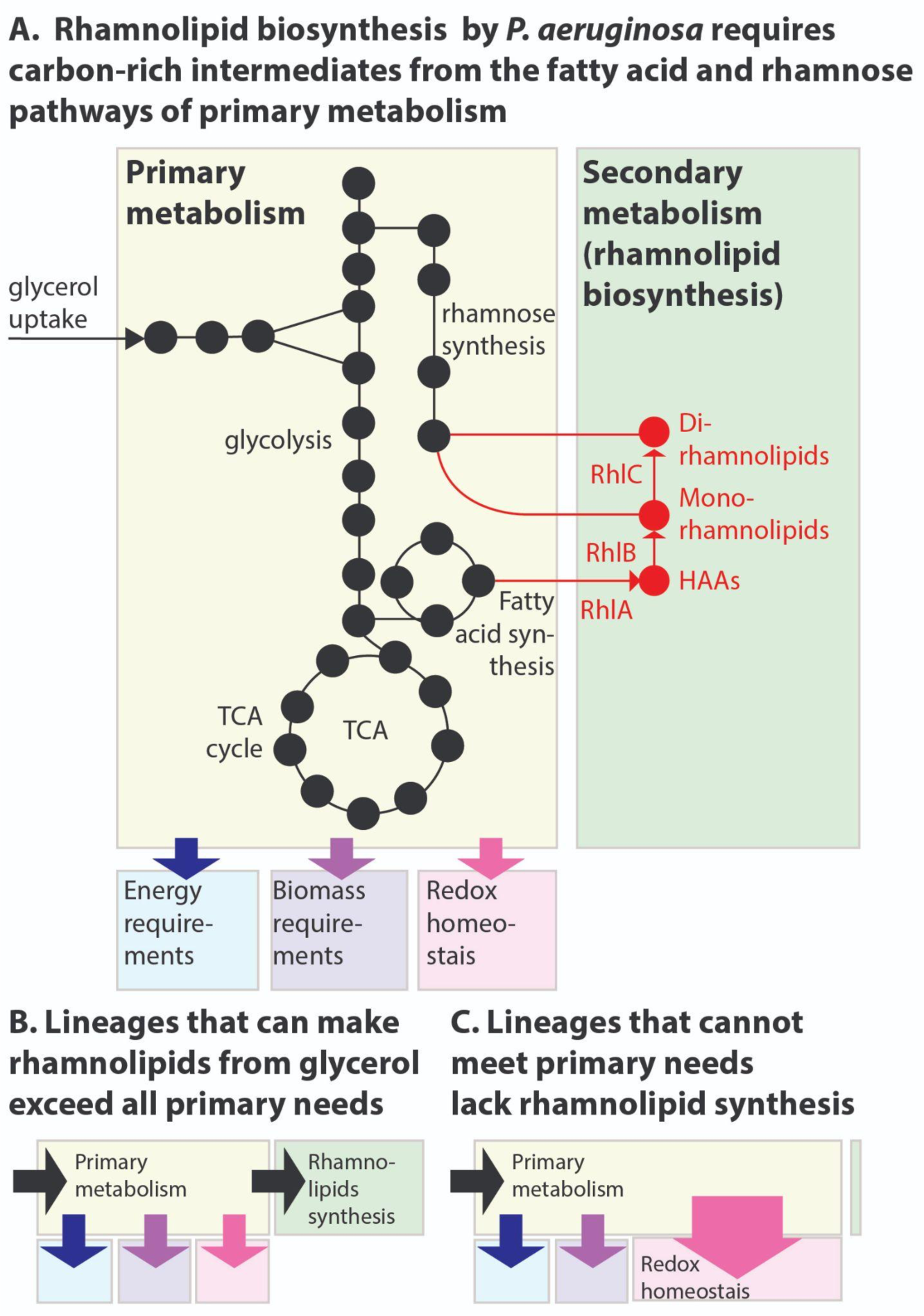
The allocation of metabolic resources made from glycerol in primary vs secondary metabolism. **A**. Growth and rhamnolipid biosynthesis on glycerol as the sole carbon source places a strong burden on *P. aeruginosa*: The bacterium has to make all the molecules needed for energy, biomass and redox homeostasis, and rhamnolipid biosynthesis—a secondary metabolic pathway—competes for those resources. **B**. *P. aeruginosa* lineages that retain the ability to make rhamnolipids from glycerol meet their primary needs and use excess resources for secondary metabolism. **C**. *P. aeruginosa* lineages that lost the ability to make rhamnolipids from glycerol were also worse at reducing the oxidative stress produced from growth in glycerol; the needs imposed on primary metabolism, such as maintaining redox homeostasis, may leave insufficient resources for secondary metabolism, explaining the loss of rhamnolipid synthesis.

What were the particular selective pressures that led to the non-producer lineages? The answer to this question remains less clear. Non-producers lacked genes (**Supplementary Table 1**) that suggested each non-producer lineage had a particular path which led to their phenotype. Importantly, though, all three genes of the secondary pathway (*rhlA*, *rhlB*, *rhlC*) remained intact, and even non-producer lineages might be able to make surfactants in conditions different from those we analyzed here. More likely, the impaired ability to reduce oxidative stress—which we linked to the inability to make rhamnolipids from glycerol—is a maladaptive consequence of specific adaptations experienced by non-producer lineages. Since the strains were isolated from hospitalized patients—a host-associated environment where we may expect stresses such as ROS imposed by the immune system in their fight against pathogens (67) and by antibiotics treatment (65)—it is unlikely that the selection was for a worse response to oxidative stress. The producer lineages could quench oxidation faster than non-producers. Our mathematical model explains this by noting that accumulation of oxidative species burdens primary metabolism and shuts down carbon overflow to rhamnolipids.

Experiments in PAO1 and clinical isolates showed previously that *P. aeruginosa* decreases expression of the key quorum-sensing genes (*lasI*, *lasR*, *rhlI*, *rhlR*, *pqsA* and *pqsR*) in response to oxidative stress caused by exposure to H_2_O_2_ (65). Considering that quorum sensing regulates hundreds of genes (68), shutting down rhamnolipid secretion could be part of a broader response to reduce oxidative stress, and saves resources for primary metabolism. But for a strain capable of reducing the stress produced by glycerol metabolism, adding an extrinsic oxidative stress induced expression of the rhamnolipid synthesis operon *rhlAB*. Lacking the Δ*rhlA* gene increased the baseline expression, but not the rate of *rhlAB* increase with H_2_O_2_, suggesting an intricate regulation. The lack of *rhlA* did not impact growth compared to the wild type even in the highest H_2_O_2_ concentrations. This is also consistent with our flux-balance analysis of *Pseudomonas* metabolism, where shutting down rhamnolipid secretion did not impact the growth rate because the excess carbon could be alternatively secreted in other forms such as acetate.

We detected nuanced consequences of deleting rhamnolipid synthesis in the strain PA14, one of the lineages capable of dealing with glycerol stress. The Δ*rhlA* isogenic mutant of PA14 had lower levels of fMet (50) which, interestingly, also happened in the non-producer lineages analyzed here. fMet plays a role in translation initiation, and in the quality control mechanisms that degrade misfolded proteins (69). *Bacillus subtilis* lacking Formyl-Methionine Transferase— the enzyme attaching a formyl group to methionine loaded on tRNA^fmet^—are more sensitive to H_2_O_2_ and defective in swarming (70). This link between fMet, oxidative stress and the loss of rhamnolipid synthesis is intriguing and warrants further research. Also the Δ*rhlA* mutant had higher levels of gamma-Glutamylcysteine (**Supplementary Fig. 8**), the immediate precursor of glutathione. Since glutathione is a well-known antioxidant that protects cells from oxidative damage (71), the result could indicate a mild oxidative stress. The Δ*rhlA* mutant expressed *rhlAB* at a higher level than the wild-type, suggesting a futile attempt to synthesize rhamnolipids. The fatty acid biosynthesis pathway that provides precursors for rhamnolipid production regenerates NAD(P)+ from NAD(P). Lack of rhamnolipid synthesis could perturb this pathway causing minor increases in oxidative stress.

The new model augments our model of rhamnolipid secondary metabolism (**Fig. 6**), helps interpret the inconsistent distribution of rhamnolipid synthesis across the *P. aeruginosa* tree, and improves our understanding the sophisticated regulation that microbes have to balance the individual costs and population benefits of secondary metabolism. There are many products of secondary metabolism that contribute to the outsize impact that microbes have on the macroscopic world. Products of one microbial species can impact other species—cooperatively or competitively—potentially amplifying microbial functions in diverse communities such as the human microbiome (59). We’ve only sampled the space of microbial secondary metabolism (7), but a general guiding principle emerges: natural selection will favor sophisticated regulation that conciliates the individual-level costs and the population-level benefits of secondary metabolism.

## Materials and Methods

### Media

Glycerol synthetic media were prepared with 800 ml of Milipore water, 200 ml of 5X minimal salts buffer, 1ml of 1 M magnesium sulphate, 0.1 ml of calcium sulphate with 0.5 gN/l nitrogen (ammonium sulphate), iron at 5 μM (iron III sulphate) and 3.0 gC/l glycerol, respectively.. 5X stock minimal salts buffer was prepared with 12 g of Na_2_HPO_4_ (anhydrous), 15 g of KH_2_PO_4_ (anhydrous) and 2.5 g of NaCl into 1 l water.

### Growth curve assays

The clinical isolates were inoculated in 3 mL of lysogen broth (LB) and incubated at 37°C overnight with shaking at 250 rpm. 500 μL of cell culture was pelleted and washed 3 times with PBS. 0.0025 OD_600_ units were inoculated into 3.0 gC/l glycerol minimal medium in BD Falcon (BD Biosciences, San Jose, CA) 96 well flat-bottom plates, with 150 μL of suspension per well. The plate was incubated during 48 hours at 37°C in a Tecan Infinite M1000 or Tecan Infinite M1000 Pro plate reader (Männedorf, Switzerland), with an orbital shaking of 4 mm of amplitude. OD_600_ was measured in 10 minutes intervals.

### Rhamnolipid production quantification

The production of rhamnolipids was assessed by drop-collapse assay (72, 73). We placed 50 μL of the culture’s supernatant on a polystyrene surface (the lid of a 96 well plate). The presence of rhamnolipids decreases the surface tension of the liquid, making the drop collapse. We considered a strain rhamnolipid non-producer if the drop does not spread, a mild-producer if spreads, but not enough to pass over the lid’s circle that corresponds to the wells, and producer if spreads over it.

### Pangenome analysis

The pangenomes of our clinical *P. aeruginosa* isolates were identified using the roary tool (74) using the output of genome annotation from prokka (75). The core genes need to have at least 95% identity using blastp and to be present in at least 99% of all genomes, and then were aligned with the MAFFT method (76).

### Phylogenetic analysis

The phylogenetic tree of clinical *P. aeruginosa* strains was constructed from core genomes as previously described (46). Moran’s I test was carried out using the ape package in R (77). The ancestral state of swarming and rhamnolipid production was reconstructed using corHMM package (78).

### Growth curve analysis

We first normalized the growth curves in different experiments (plates) by matching their corresponding controls (PA14). The plate-specific coefficients were obtained by solving a linear regression model ‘log(OD of PA14) ∼ Plate + Time’ (the Wilkinson notation) using Matlab function fitlm, where Plate is plate ids and Time is time points (both are categorical variables). Then the growth curves in each plate were normalized by multiplying the adjustment coefficient of the plate. These normalized growth curves were then smoothed by Savitzky-Golay filter (sgolayfilt function in Matlab). The growth phases (phase I, II, III) were determined by the following rules: (phase I): first-order derivative >0 and second-order derivative >0; (phase II) first-order derivative >0 and second-order derivative <0; (phase III) first-order derivative <0. The mean and maximum specific growth rate for each growth phase were estimated from the best-fit model (highest R2) among three models of bacterial growth curves: (1) exponential model *y* = *y*_0_ + *y*_1_*e*^μ*t*^; (2) Zwietering-Logistic model 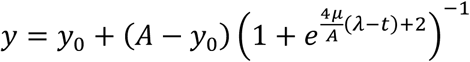; (3) Zwietering-Gompertz model 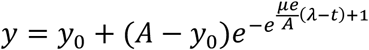, where *y*_0_, *y*_1_, *μ, λ, A* are the parameters to fit. Both non-negative factorization and random forest regression were performed in python, by sklearn.decomposition.NMF and sklearn.ensemble.RandomForestClassifier respectively.

### Metabolite extraction

All *P. aeruginosa* strains were grown until the end of exponential phase of growth in glycerol minimal medium. Bacteria was then loaded into 0.25 μm nylon membranes (Millipore) using vacuum, transferred to pre-warmed hard agar plates with the same medium composition and incubated at 37°C during 2.5 h. The filters were then passed to 35 mm polystyrene dishes (Falcon) with 1 mL of 2:2:1 methanol:acetonitrile:H_2_O quenching buffer and incubated there during 15 minutes on dry ice. Cells were removed by scraping and the lysate containing quenching buffer was transferred to 1.5 mL tubes and centrifuged at 16000 rpm for 10 minutes at 4°C. Supernatant transferred to fresh tubes and stored at -80°C.

### Metabolomic data preprocessing

The extracts were profiled using liquid-chromatography coupled to mass spectrometry (LC-MS), identifying a total of 92 compounds (Supplementary Fig. 5). Some compounds contained missing values. These missing values in metabolite abundance can be (1) truly missing; (2) present in a sample but its level is below detection limit; (3) present in a sample at a level above the detection limit but missing due to failure of algorithms in data processing. Here we assume that a metabolite with missing values in all three replicates is truly missing in the sample and removed from our analysis (Supplementary Fig. 5). However, if the missing values were only found in one or two replicates, the missing values were imputed by the average of the non-missing values. After that imputation all compounds with missing values were removed (Supplementary Fig. 5).

The peak areas were normalized using Cross-Contribution Compensating Multiple Standard Normalization (CCMN) (79) with NormalizeMets R package (80). This method relies on the use of multiple internal standards. Since LC-MS lacks such internal standards, we used instead a set of metabolites assumed to be constant across all the strains. They were selected with a Kuskal-Wallis test, adjusting the *p-*value with Benjamini-Hochberg method. The ones with a *p-*value above 0.05 were considered constant (pyruvate, methylglyoxal, (S)-2-Acetolactate, Tyramine, D-Glucose, (S)-Lactate, N-acetyl-L-glutamate 5-semialdehyde, 4-Aminobutyraldehyde and Glycine), therefore after the normalization step they were removed (indicated in red, **Supplementary Fig. 5A**). The processed area peaks for all metabolites are included in **Supplementary Table 5**.

### Unsupervised analysis of metabolomic data

The hierarchical clustering of the normalized metabolomic data was performed using gplots R package (81), with Euclidean distance and Ward’s aggregation method. The clustering was performed with all metabolites; however, 16 metabolites (indicated in black, **Supplementary Figure 4**) with unknown compound identity were removed from **Supplementary Figure 5** and downstream analyses. These unknown compounds are not artifacts arising from LC-MS instruments since their peaks were detected in all strains. Fumarate and guanosine, which were only putative initially, were bioinformatically confirmed as all other compounds with the same molecular weight are produced by enzymatic reactions missing in our clinical isolates. The Principal Component Analysis was done with R’s base method, using also the normalized data and previously removing the 16 metabolites with unknown identity. The plot was obtained using ggplot2 R package (82).

### Metabolic pathway enrichment

The differential metabolites between rhamnolipid producers and non-producers was determined by a Mann-Whitney test, with *p*-values adjusted with Benjamini-Hochberg method and a significance level of 0.05. These compounds were fed to FELLA algorithm (49, 83). FELLA retrieves a graph describing the relationships among pathway, module, enzyme, reaction and compounds of *Pseudomonas aeruginosa* strain UCBPP-PA14 from the KEGG database. The graph was then used as an input network of differential compounds for its diffusion algorithms (84). The output of FELLA consists of all subnetworks of the entries predicted to have a high connectivity with the differential compounds. We filtered the subnetwork entries to keep only metabolic pathways. The entries shown in the supplementary table 3 are the ones with a significant probability of receiving part of the simulated flux.

### OPLS-DA model

OPLS-DA model of metabolomics data was built using ropls R package (85), fixing the number of orthogonal components to 3. R^2^ and Q^2^, key parameters for assessing the validity of the model, were assessed with 7-fold cross validation. The significance of the model was determined by permutation test (n = 2000). The *p*-value corresponds to the proportion of Q^2^perm above Q^2^. With a *p-*value below 0.05 we considered the model significant. The loadings of the predictive component of the model were extracted to determine how each metabolite contributes to the separation according to the phenotype.

### Genome-scale modeling

Custom Python codes were developed with the COBRApy package (86) to carry out all metabolic flux modeling and simulations in the paper. Since iJN1411 model was developed for *Pseudomonas putida*, we removed genes and associated reactions that are missing in all our strains but present in the iJN1411 model. The futile cycles involving NADH, NADPH, and GSH were also removed. The modified iJN1411 model was further expanded by adding rhamnolipid biosynthesis pathway involving 9 new metabolites and 12 new reactions.

The boundary fluxes of the model were set to mimic the composition of the glycerol minimum medium. For C:N=10, the lower bounds of glycerol and ammonium fluxes were set to -10 and -3 respectively. For C:N=3, the lower bounds were set to -3 and -3 respectively. The flux unit is mmol/gDW/h throughout the paper. To constrain the total producing flux of NADH (the same for NADPH and GSH) at a certain value *C*, we first defined a binary variable *X_k_* for each NADH-involving reaction *k* to indicate whether NADH is produced by this reaction. Given the stoichiometric coefficient of NADH in this reaction (*s_k_*) and its flux value (*f_k_*), the mathematical constraints for *X_k_* was set by *X_k_* = 1 for *s_k_f_k_* > 0 and *X_k_* = 0 otherwise. Therefore, the constraint that equalizes the total NADH producing flux and a constant *C* is simply ∑*_k_ s_k_X_k_f_k_* = *C*. However, both *X_k_* and *f_k_* are variables and such quadratic constraint has not yet been supported by COBRApy. We overcame this difficulty by defining *A_k_* = *X_k_f_k_* and linearized the product with the following two inequalities: *X_k_l_b_* ≤ *A_k_* ≤ *X_k_u_b_* and *f_k_* − (1 − *X_k_*)*u_b_* ≤ *A_k_* ≤ *f_k_* − (1 − *X_k_*)*l_b_*, where *l_b_* and *u_b_* are the lower and upper bounds of *f_k_*. The two constraints ensure that *A_k_* = 0 when *X_k_* = 0 and *A_k_* = *f_k_* when *X_k_* = 1. The minimum/maximum flux values of byproduct secretion were simulated by flux variability analysis at maximum growth rate.

### Detection of hydrogen peroxide (H_2_O_2_)

The H_2_O_2_ level in the extracellular medium was quantified with Amplex® Red Hydrogen Peroxide/Peroxidase Assay Kit (Invitrogen, Carlsbad, USA, Catalog no. A22188). 500 μL of cell suspension were spinned down after overnight growth in LB medium, washed twice in PBS and normalized to OD 1. Each reaction was done in a final volume of 100 μL, with a final concentration of 50 μM of Amplex® Red reagent, 0.1 U/mL of HRP (Horseradish Peroxidase) and 0.2 cell OD, in glycerol synthetic medium, in BD Falcon (BD Biosciences, San Jose, CA) 96 well flat-bottom plates. The first column of the plate corresponded to the reaction without cells (the volume was substituted by PBS), and the last column instead of cells contained H_2_O_2_ (final concentration 10 μM). OD_600_ and fluorescence 530/590 nm was measured in 10 minutes intervals (48 h 37°C).

### RNA-seq and data process

The *P. aeruginosa* strains were inoculated into LB and grew with agitation for 16h. The overnight culture was diluted in glycerol synthetic medium the next day and grew to late exponential phase to harvest RNA. Cells were harvested by centrifuge and then immediately resuspended in RNAprotect (QIAGEN). The RNA was extracted using the RNeasy Mini Kit from QIAGEN. The standard RNA sequencing was done by Genewiz (Genewiz, Inc; South Plainfield, NJ) using pair-end indices of 150bp read length on an Illumina HiSeq platform. The raw fastq files first went through quality check by fastQC (87), and aligned with STAR (88). The final output was generated using DEseq2 (89) package in R. The read counts for each gene is included in **Supplementary Table 6**. The SRA accession number for the RNAseq data is listed in Supplementary Table 7.

### RLQ analysis

To test for the functional associations between genes and rhamnolipid production, we adopted the RLQ analysis (ade4 package in R) that was developed for identifying the associations between traits and environmental variables. Here we applied it to RNAseq analysis to understand the main co-structures between gene functions and phenotypes mediated gene expression. The theory of RLQ analysis is described in detail elsewhere (90). Briefly, it builds on the simultaneous ordination of three tables: a R table (categorical) describing rhamnolipid production (strong-, mild- and non-producers) for 11 selected isolates (each replicate was treated as an independent observation), a L table (continuous) describing the RNAseq counts of 1,474 core genes shared among the 11 isolates, and a Q table (binary) describing 120 KEGG pathways for the 1,474 genes. We first applied corresponding analysis (CA) to table L and Q and principal component analysis (PCA) to table R. In the CA of R and Q, each strain and gene were weighted by their total RNA expression levels respectively. The three separate ordinations are then used as inputs to the rlq function. The output of the function includes coefficients for the gene functions and phenotypes (loadings) as well as the scores of strains (out of phenotypes) and scores of genes (out of gene functions). RLQ analysis maximizes the squared cross-covariances between the two sets of scores weighted by gene expressions.

### Hydrogen peroxide (H_2_O_2_) degradation

H_2_O_2_ was removed from environment by cells which degrade H_2_O_2_ intracellularly. The cumulative H_2_O_2_ removal curve was calculated by subtracting the values of emission of the wells containing each strain from the values of emission of the wells without cells or H_2_O_2_. Then they were normalized to the relative scale by dividing their values by the corresponding values (at the same time points) of the control strain (PA14) in the same experiment (plate). These normalized curves were then smoothed by the Savitzky-Golay filter (sgolayfilt function in Matlab). The H_2_O_2_ removal rate per cell was calculated by dividing the first-order derivative of the cumulative removal curves by OD values. To estimate the effect of rhamnolipid production on per-cell H_2_O_2_ removal rate, we developed a linear mixed-effect model that extends the simple linear regression model by considering both fixed and random effects. The model was specified by the Wilkinson notation ‘RemovalRate ∼ Time + RHL + (1|CurveID)’, where RemovalRate is per-cell H_2_O_2_ removal rate, Time is time point (categorical variable), RHL is categorical rhamnolipid production (non-producer, mild-producer, and strong-producer), and CurveID is the unique ID for each strain and replicate. The expression ‘(1|CurveID)’ indicates that CurveID was modeled as random effects. The model was solved by fitlme in Matlab.

### Promoter activity and growth rate comparison

*P. aeruginosa* strains were inoculated into LB and overnight cultures were harvested and washed with PBS three times. Cells were diluted to OD_600_=0.1 in synthetic media and monitored using TECAN plate reader for both OD_600_ and GFP. Upon cells entering the exponential phase, H_2_O_2_ was added to the cells, which continued to be monitored over 48h. After identifying growth phase II, the promoter activity was calculated as: 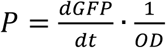. The ratio of promoter activity in each experiment was extracted as 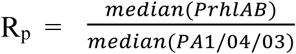. Similarly, the growth rate was calculated as 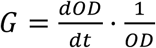, and the ratio of strains with different promoters was calculated as 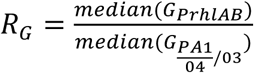. The effect of H_2_O_2_ was estimated by fitlme in Matlab as: log(R)∼a_0_*β* ⋅[H_2_O_2_]+*ϵ*, where R is the measured ratio of either promoter activity or growth rate, and *ϵ* is denoted as the random effect of experiments.

## Data availability

All experimental and simulation data that support the conclusions of this study are available from the main text and Supplementary Information. The source codes for generating the figures of this study are available from: https://github.com/guisantagui/code_PA_paper; https://github.com/liaochen1988/Source_code_for_Pseudomonas_Metabolomics_Paper; https://github.com/Jinyuan1998/PA_metabolomics_rhamnolipids_SourceCode.

## Acknowledgements

This work was supported by NIH grants U01 AI124275 and R01 AI137269-01 to J.B.X. F.R.P. and G.S. were partially supported by UIDB/04046/2020 and UIDP/04046/2020 Centre grants from FCT, Portugal (to BioISI) and by the LungCARD project (Grant Agreement n°: 734790, H2020-MSCA-RISE-2016). G.S. is recipient of a fellowship from BioSys PhD programme PD65-2012 (Ref SFRH/BD/142899/2018) from FCT (Portugal). The funders had no role in study design, data collection and analysis, decision to publish, or preparation of the manuscript.

## Supplementary Figures

**Supplementary Figure 1.**
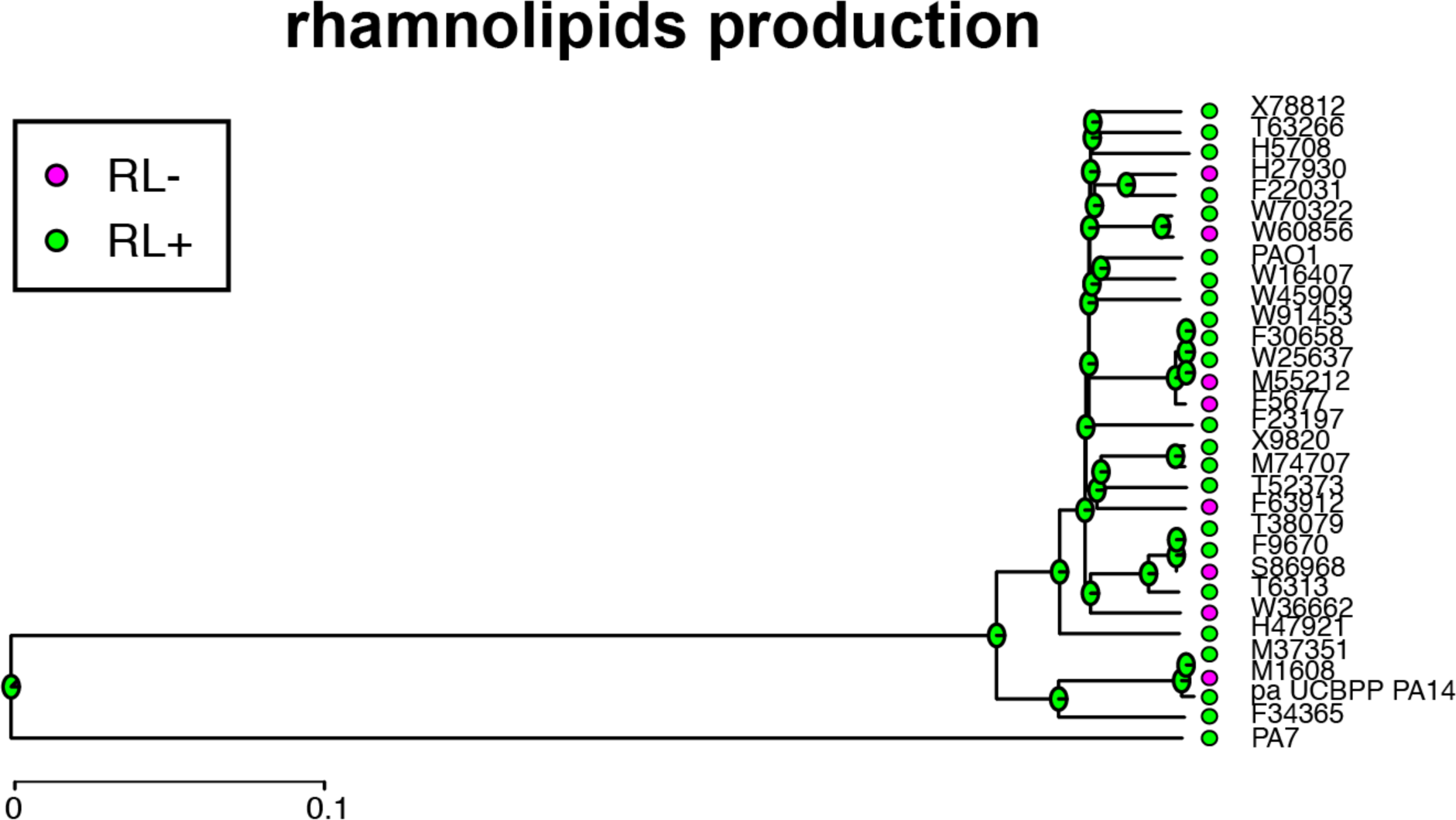
Phylogenetic ancestor state reconstruction of rhamnolipid production. Pie charts at the ancestor nodes of branches represent the relative likelihood proportion of each possible phenotypic state. We rooted the tree with PA7, a *P. aeruginosa* isolate that is often used as an outlier to root phylogenetic trees. RL, rhamnolipid production.

**Supplementary Figure 2.**
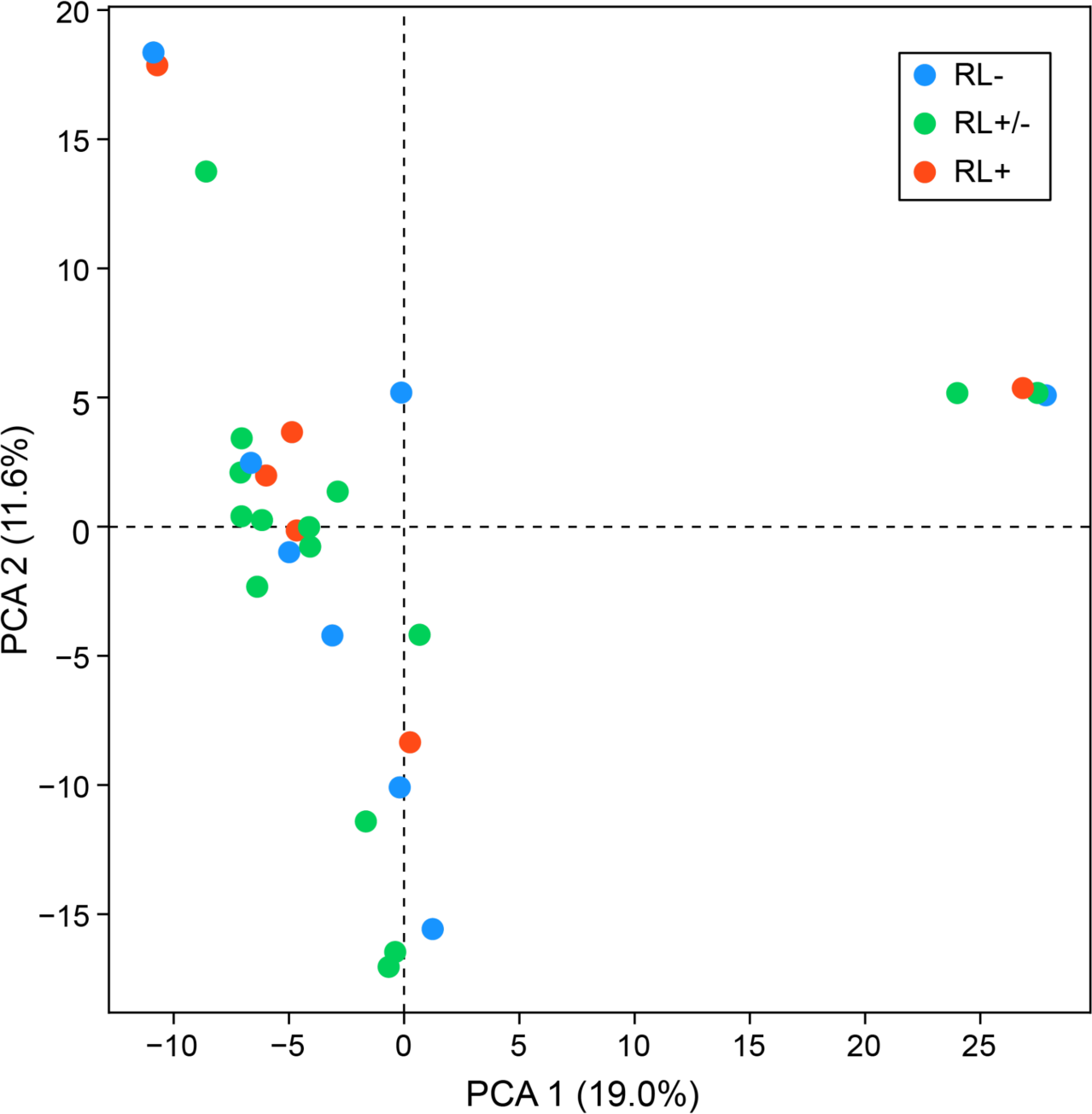
Principal component analysis (PCA) of the presence/absence of the accessory genomes.

**Supplementary Figure 3.**
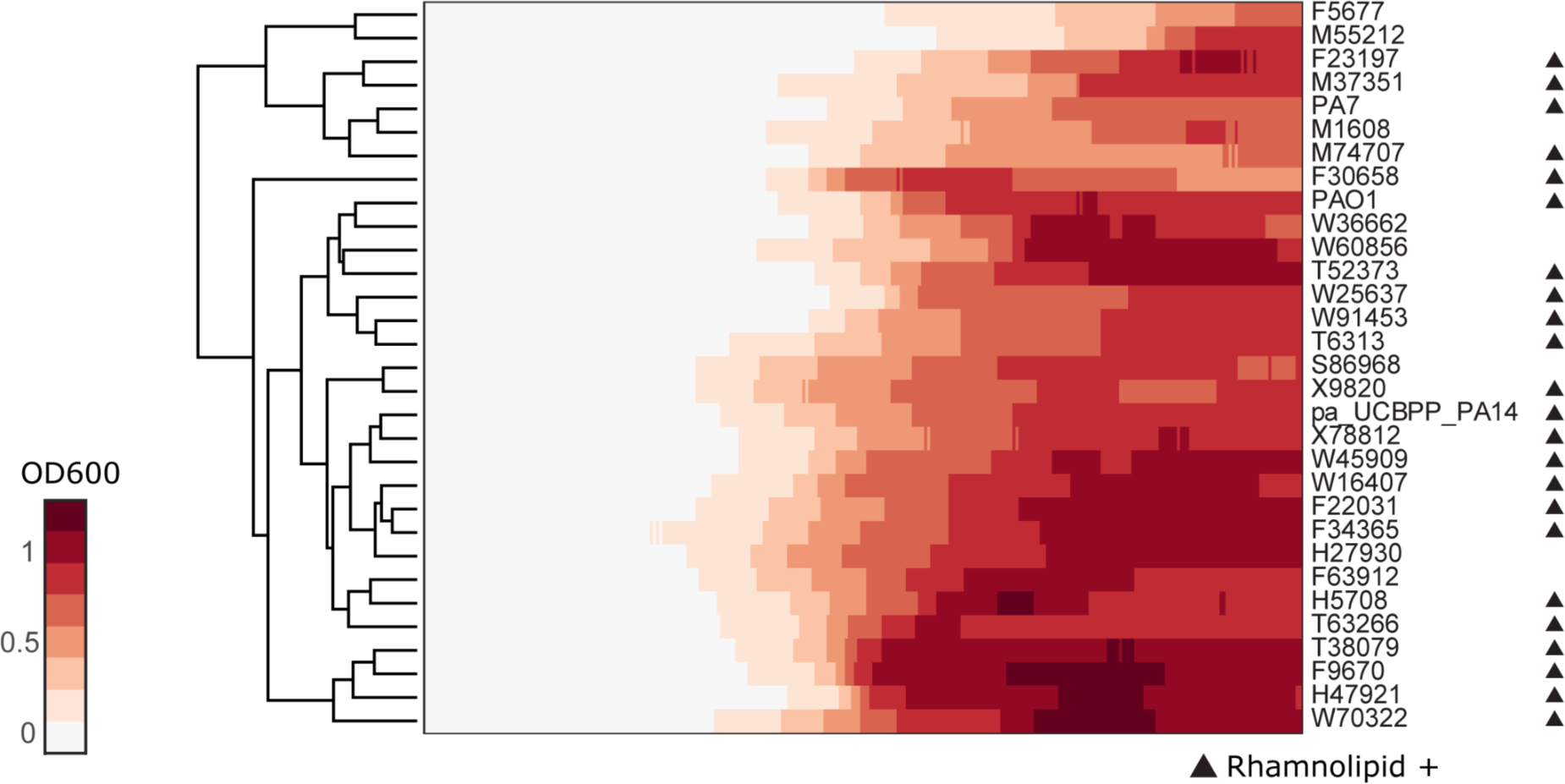
Clustergram of growth curves of *P. aeruginosa* clinical isolates and three type strains PA14, PAO1 and PA7 in glycerol minimal medium. Euclidean distance was used as the measure of similarity.

**Supplementary Figure 4.**
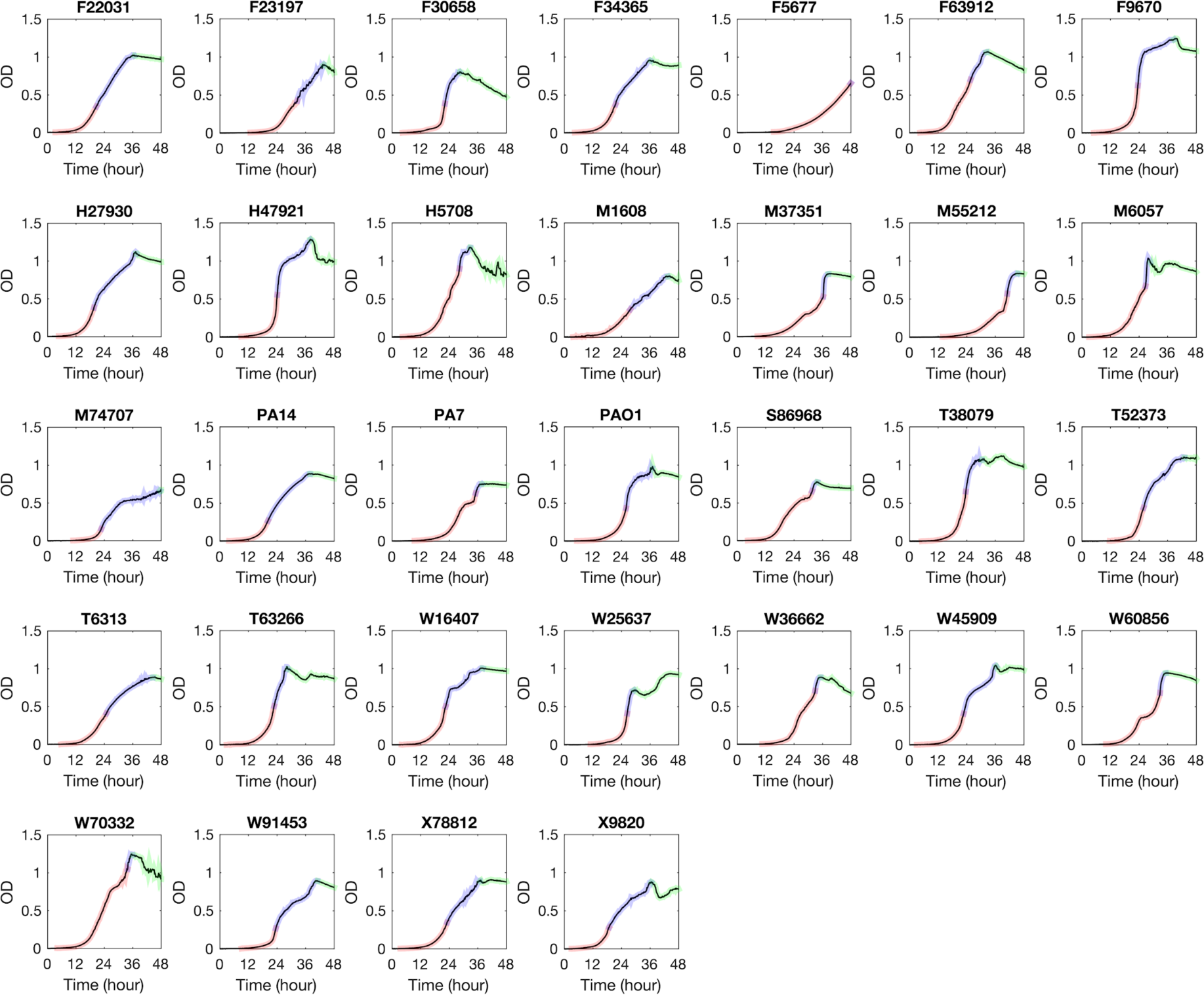
Growth curve of *P. aeruginosa* strains in glycerol minimal medium. Phase I, II, and III are colored by red, blue and green respectively.

**Supplementary Figure 5.**
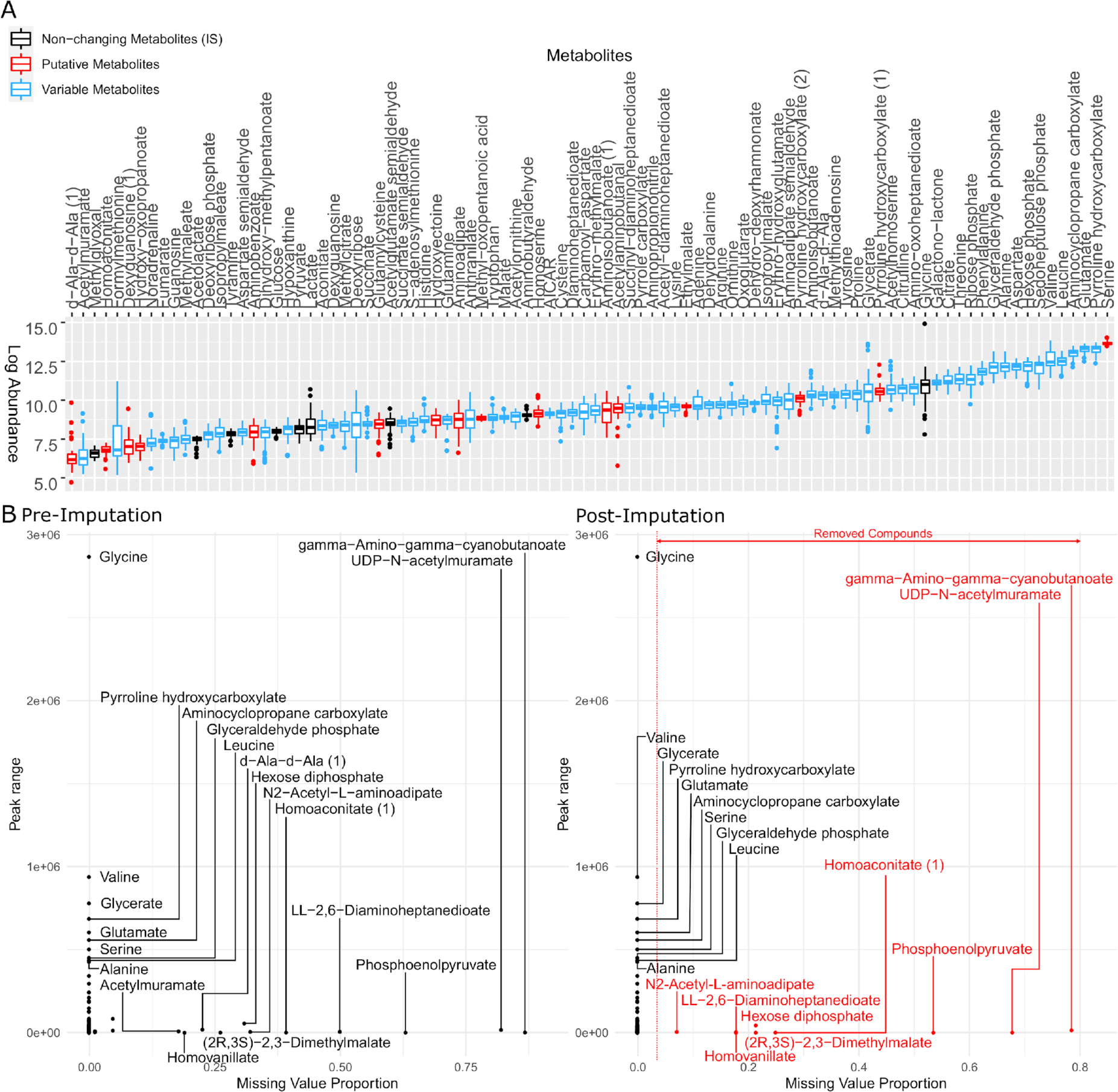
**A.** Logarithm of raw peak area of all metabolites identified by LC-MS (Liquid chromatography-Mass spectrometry). The metabolites in black were used as internal standard in the data normalization step but removed from the dataset afterwards. The metabolites in red are putative metabolites that were initially included in clustering metabolomics but later removed from the heatmap (**Supplementary Figure 6**). The metabolites in blue are variable among clinical isolates and therefore used for later on analysis. **B.** Metabolite peak area before (left panel) and after (right panel) imputation. The missing value proportion of a metabolite in the x-axis represents the frequency of missing values of the metabolite across all replicates of all strains in our study. The peak range of a metabolite in the y-axis is defined as the maximum peak area minus the minimum peak area among all non-missing values of the same metabolite across all our strains.

**Supplementary Figure 6.**
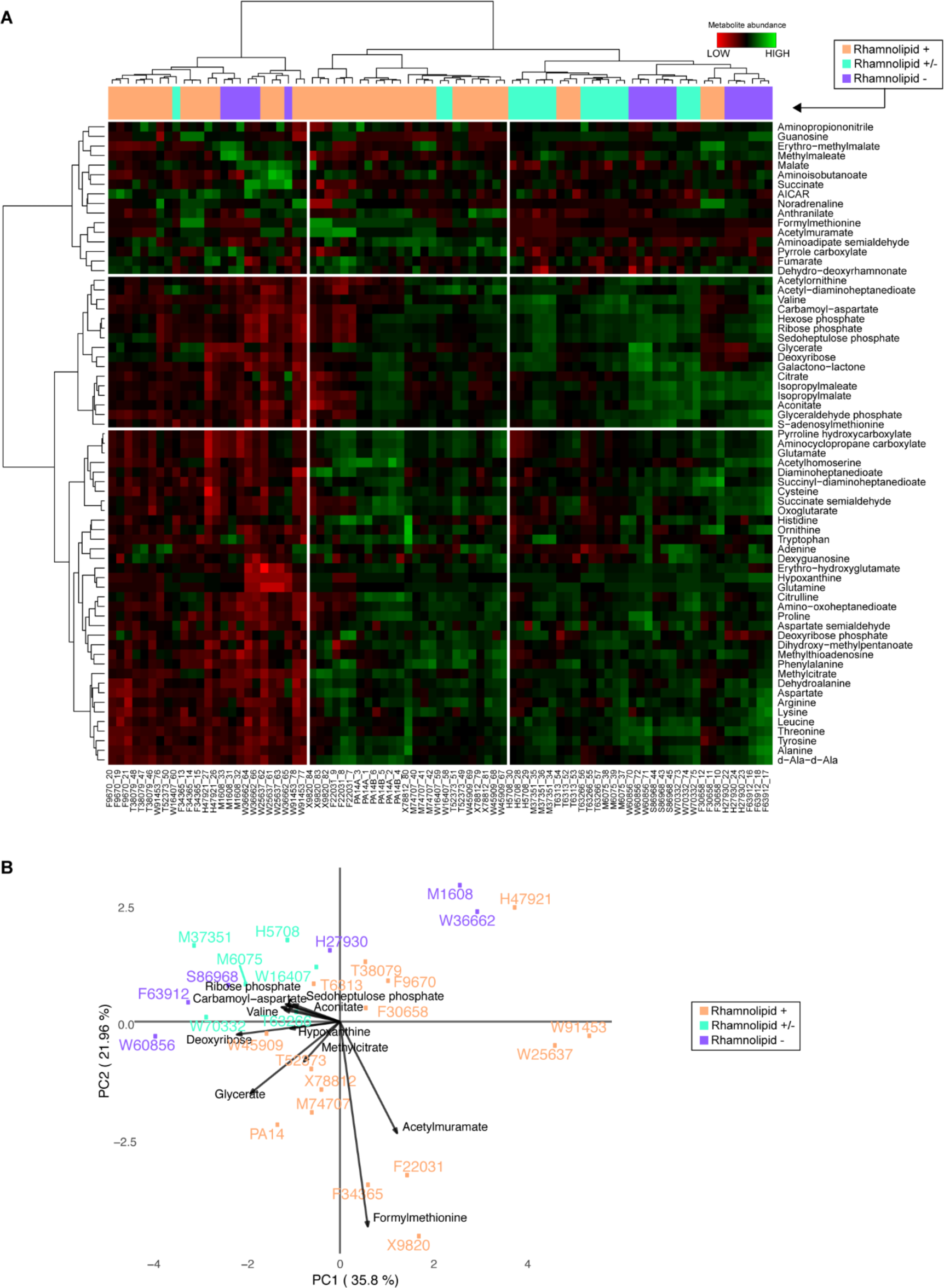
Variance and clustering analysis of metabolomics across *Pseudomonas aeruginosa* strains. **A.** Hierarchical clustering of the metabolic profiles using Euclidean distance and Ward aggregation method. Each row represents one metabolite and each column represents a specific sample (both strain and sample number are indicated). **B.** PCA plot of metabolites from LC-MS. The rhamnolipid strong producer (+) occupies different territory but the separation among the three groups can not be explained using PC1 or PC2. Strains from the purple (-) and the green (+/-) groups overlap.

**Supplementary Figure 7.**
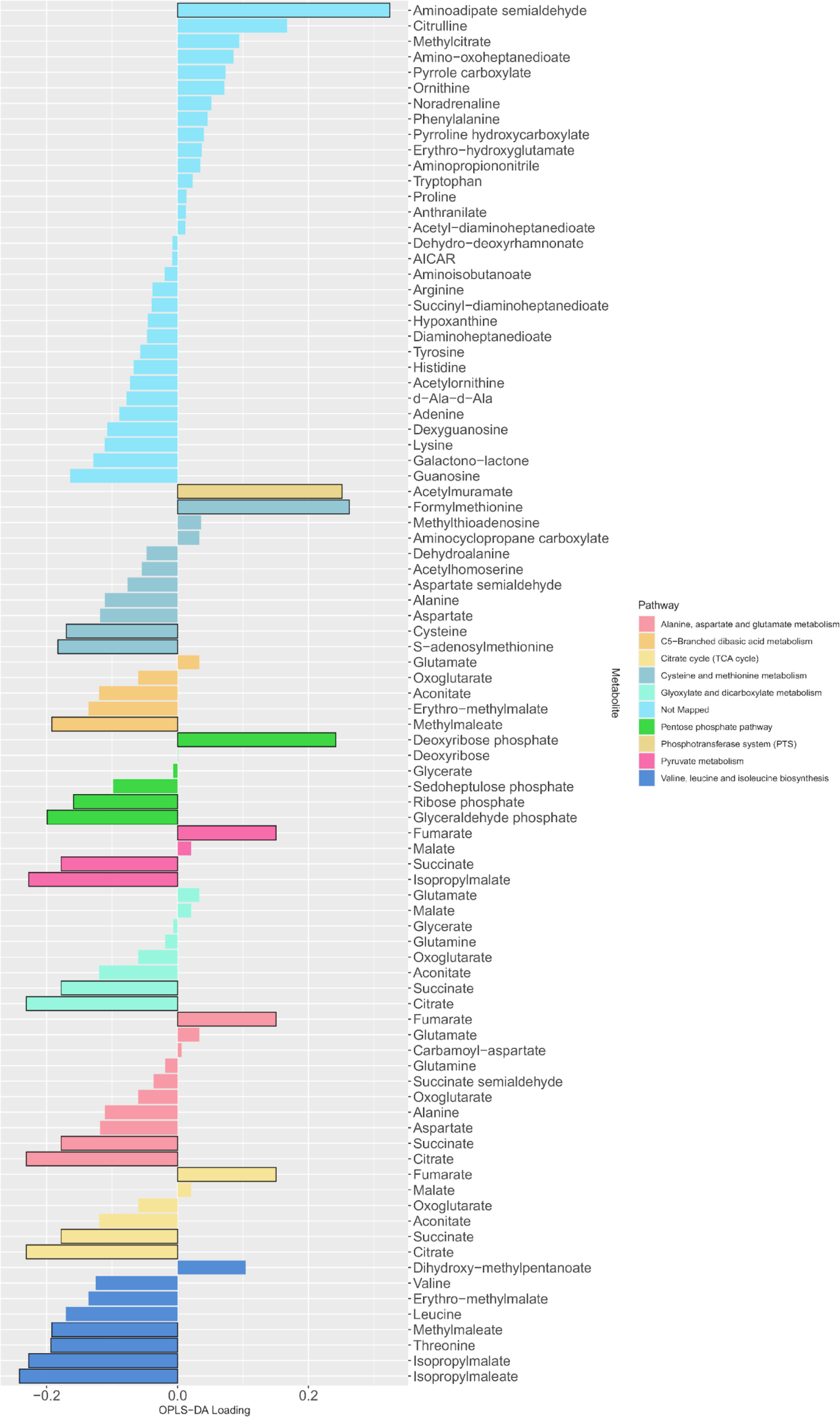
The loading values of all predictive metabolites of the OPLS-DA model. The differential metabolites between producers and non-producers were determined by a Mann Whitney test (adjusted *p-*values with Benjamini-Hochberg method) with a level of significance of 0.05 (bars with black outline) and used as input for a metabolic pathway enrichment with FELLA algorithm. The colors indicate the mapped metabolite pathway for each metabolite.

**Supplementary Figure 8.**
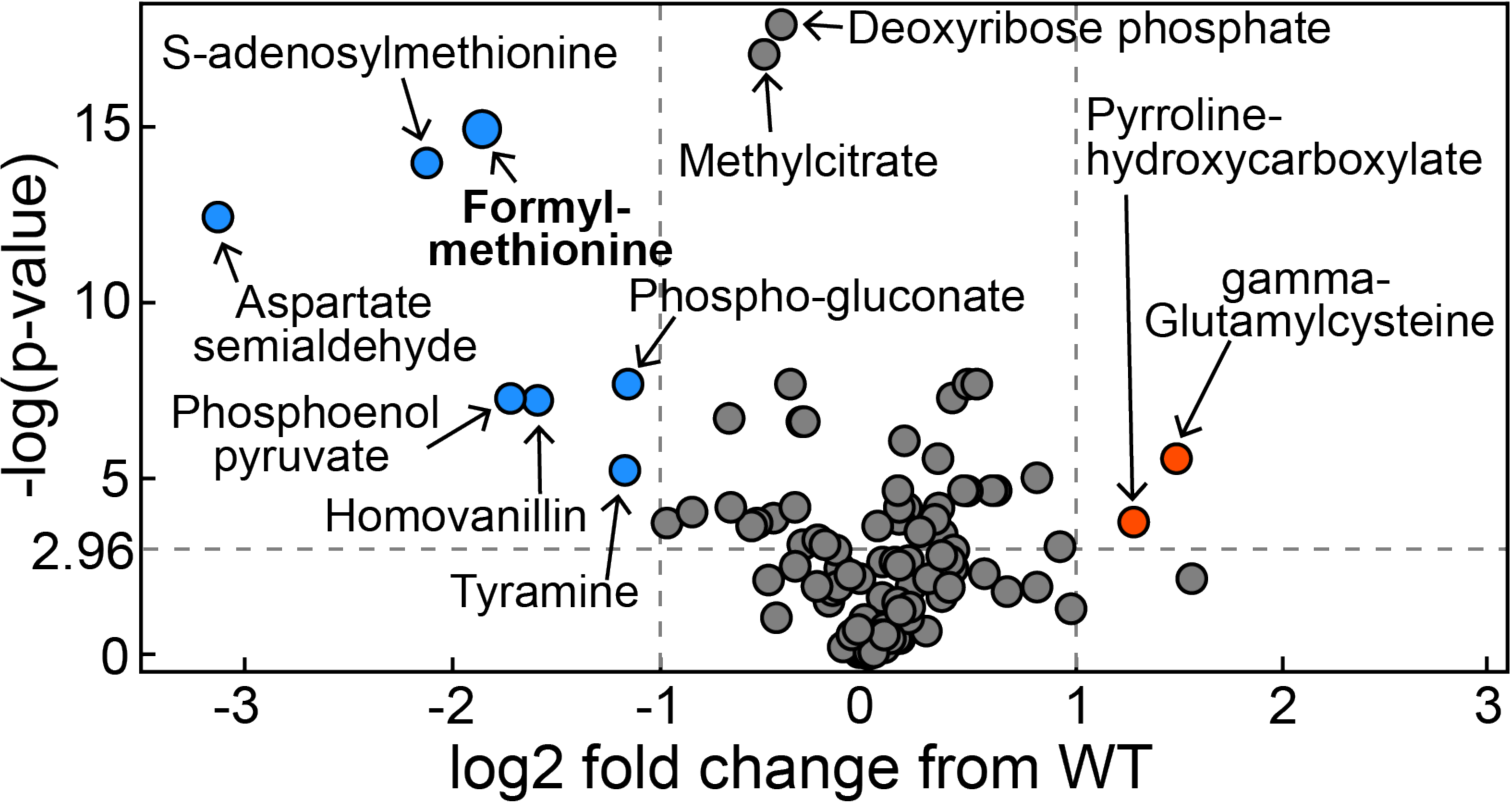
Volcano plot of metabolomics data between wild-type *P. aeruginosa* UCBPP-PA14 strain and its *rhlA* mutant grown in glycerol minimal medium (replotted with permission from (20)).

**Supplementary Figure 9.**
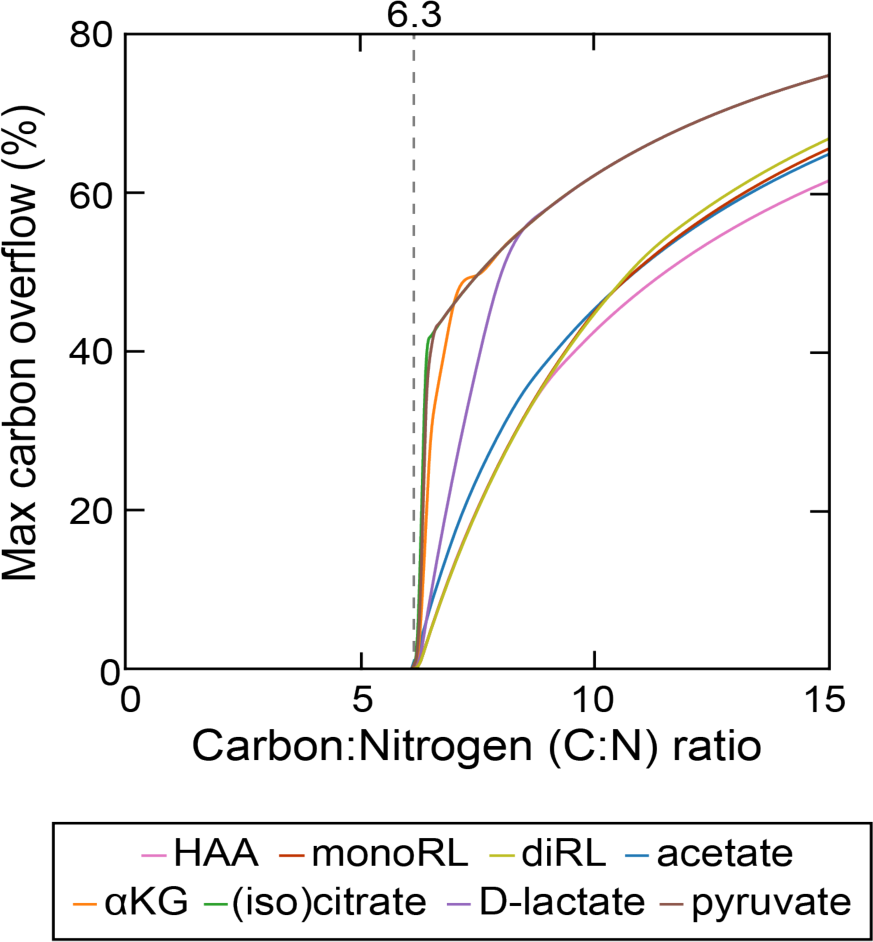
Theoretical estimation of threshold carbon (glycerol):nitrogen (ammonium) ratio above which carbon is in excess in the sense that carbon release through rhamnolipids and central carbon metabolites does not compromise biomass production. Abbreviations: HAA: 3-(3-hydroxyalkanoyloxy)alkanoate; monoRL: monorhamnolipid; diRL: dirhamnolipid; aKG: alpha-ketoglutarate.

**Supplementary Figure 10.**
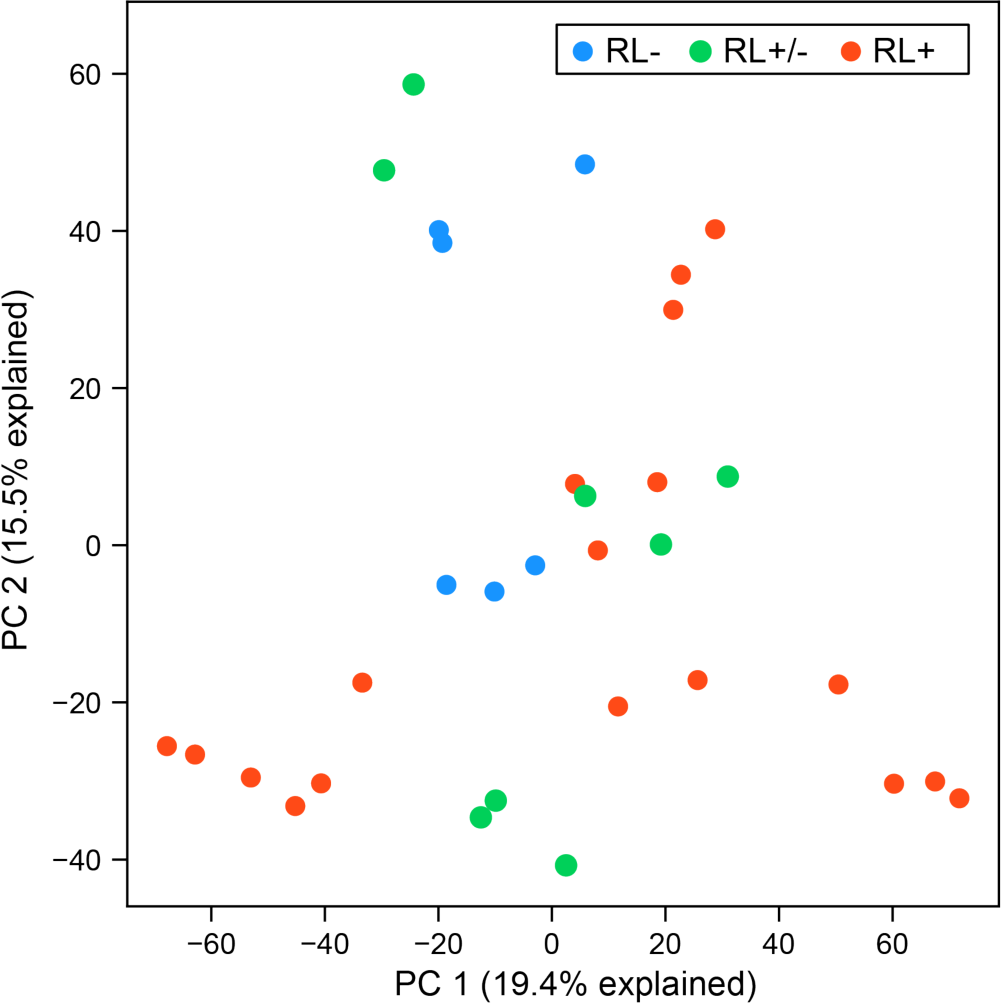
PCA plot of RNA expression colored by rhamnolipid production phenotypes. Each dot represents a replicate of an RNA-seq experiment. PC: principal component; RL-: rhamnolipid non-producer; RL+/-: mild rhamnolipid producer; RL+: strong rhamnolipid producer.

**Supplementary Figure 11.**
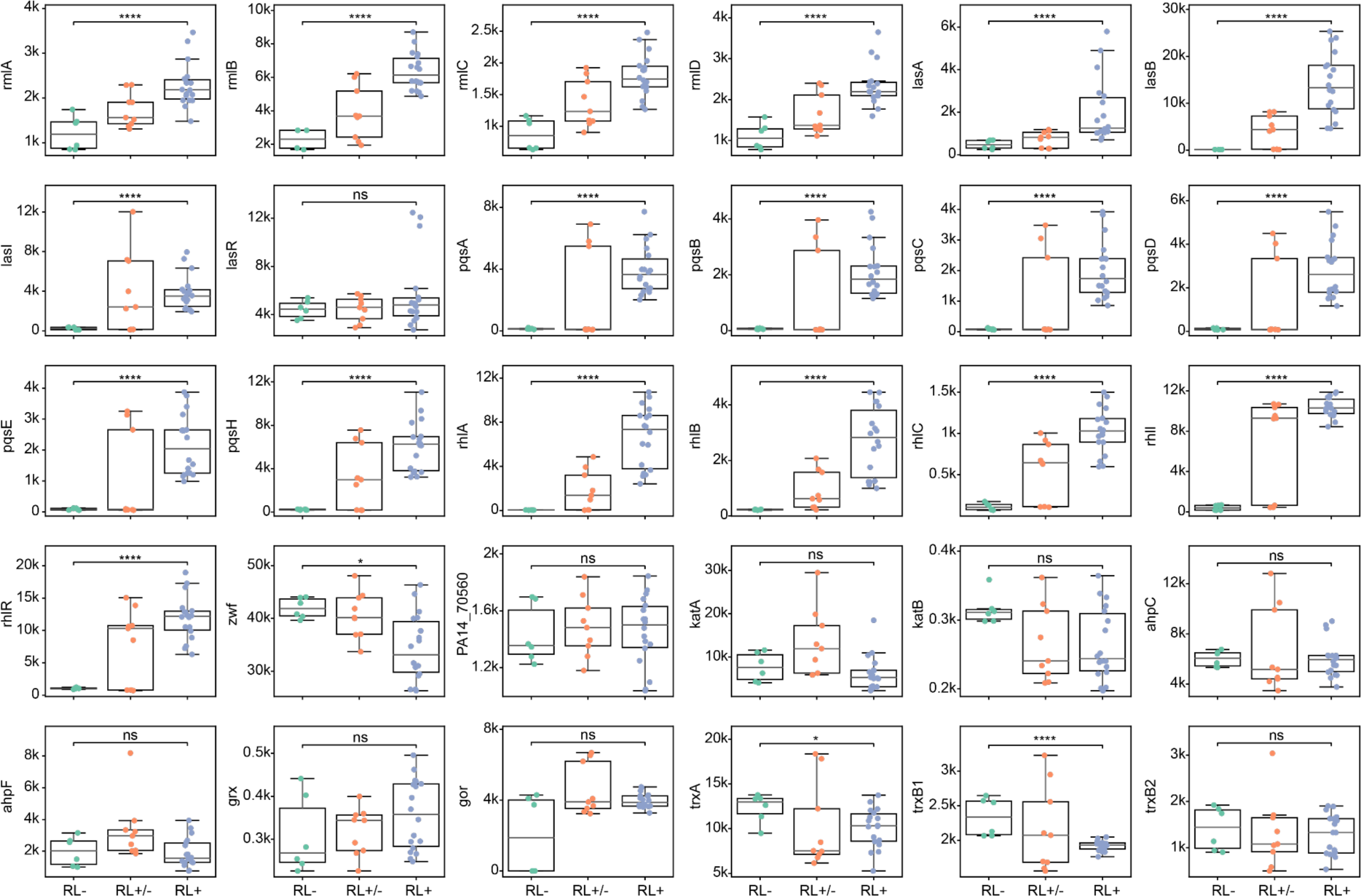
Comparisons of expression for selected genes across strains with different abilities of rhamnolipid biosynthesis. Each dot represents a replicate of an RNA-seq experiment. RL-: rhamnolipid non-producer; RL+/-: mild rhamnolipid producer; RL+: strong rhamnolipid producer. Mann-Whitney U test with FDR method for multitest correction. ****, P<0.0001; *, P<0.05. ns, non-significant.

**Supplementary Figure 12.**
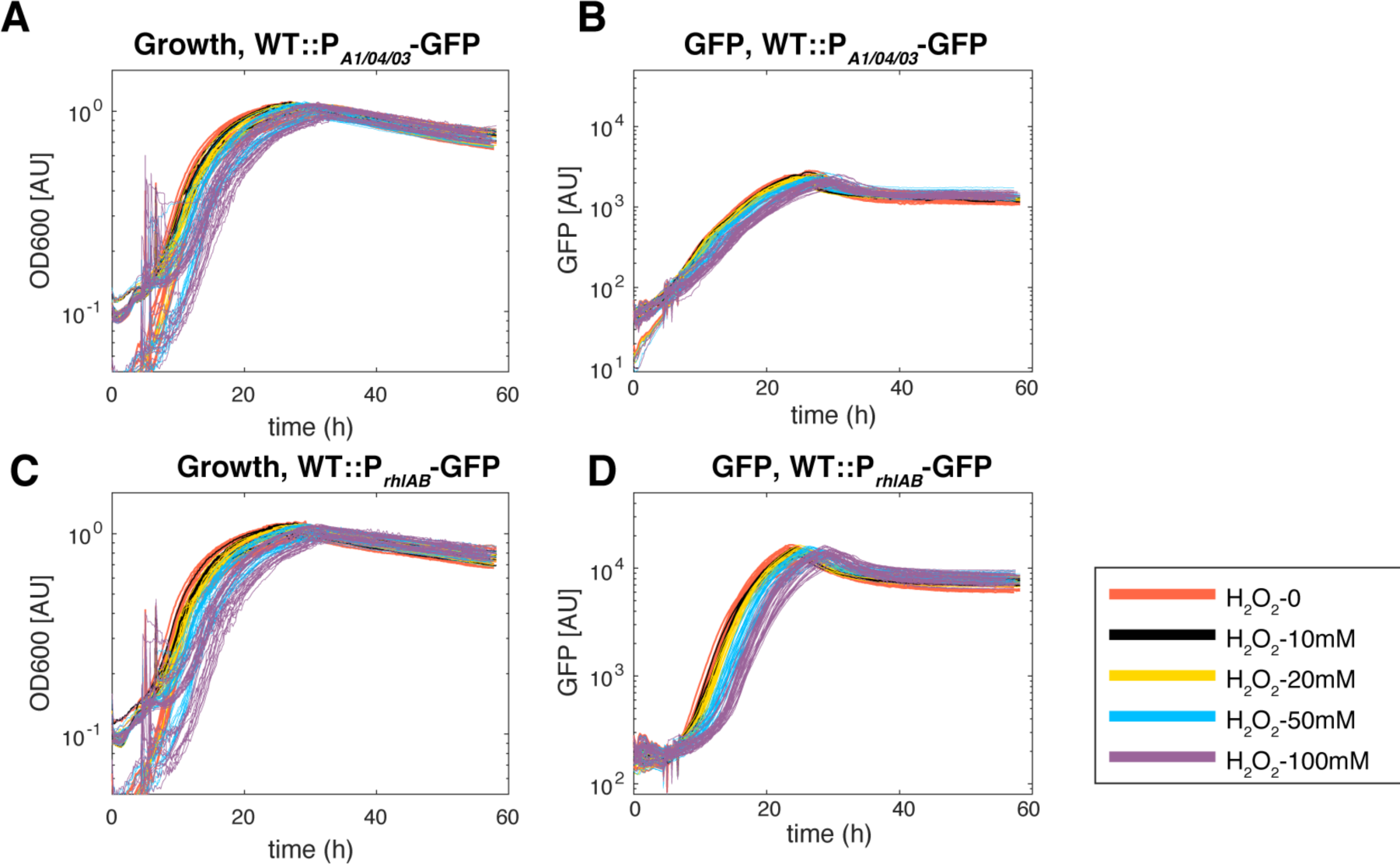
Cell growth and *rhlAB* expression under extra oxidative stress in glycerol minimal media with wild type strain. Each line represents the data from a single well. The disturbance of the lines were caused by added H_2_O_2_ to the media.

**Supplementary Figure 13.**
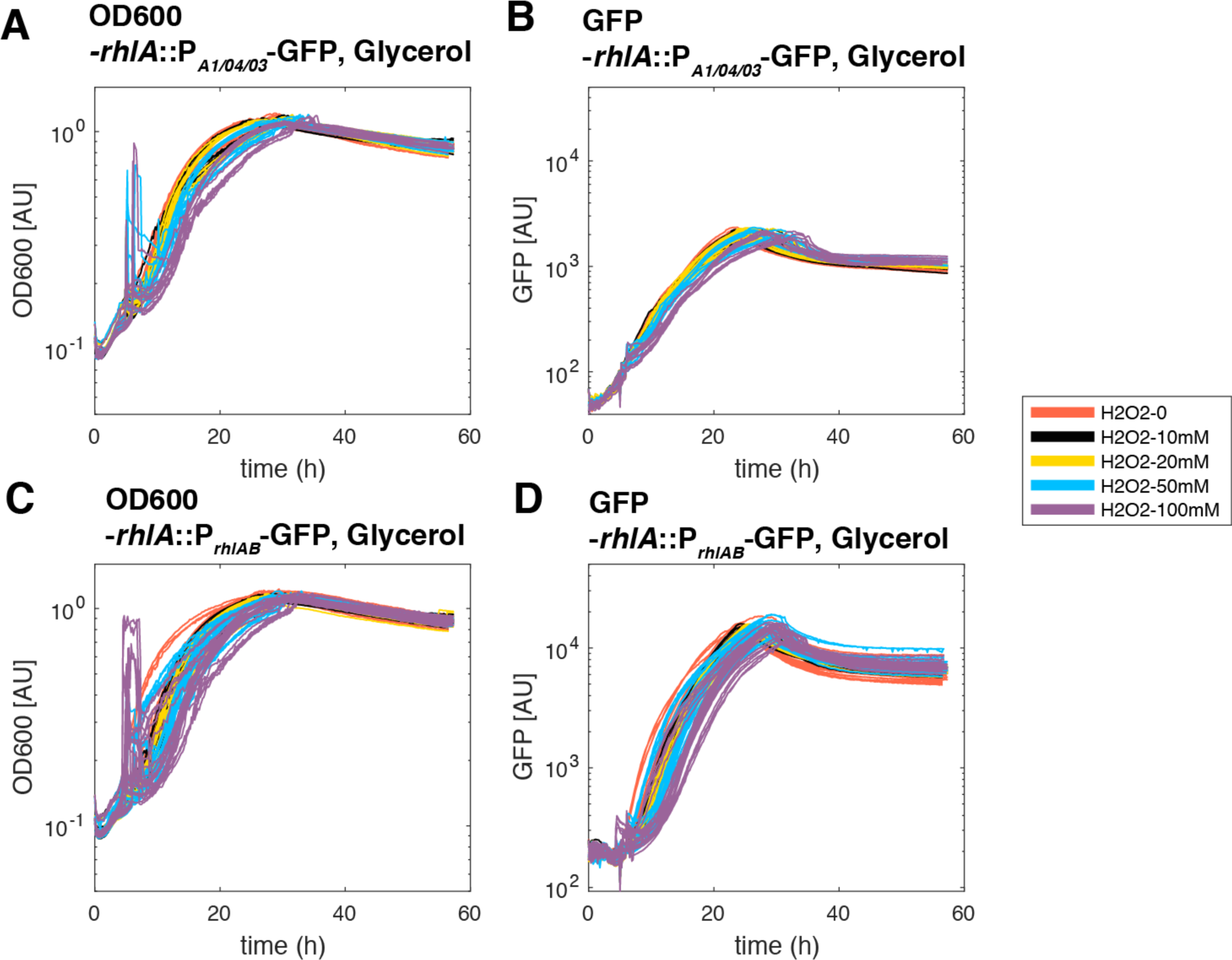
Cell growth and *rhlAB* expression under extra oxidative stress in glycerol minimal media with *ΔrhlA* strain. Each line represents the data from a single well. The disturbance of the lines were caused by added H_2_O_2_ to the media.

## Supplementary Data

**Supplementary Table 1. Presence and absence of genes across the genomes of our clinical isolates**

**Supplementary Table 2. Quantitative values for the 7 local features of phase I, II and III (phase start time point, phase duration, phase initial OD, OD change, area under the curve, mean specific growth rate, maximum specific growth rate) of growth curves.**

**Supplementary Table 3. Different pathways that are significantly different in rhamnolipid producers and non-producers identified by FELLA.**

**Supplementary Table 4. Correlations between gene expressions or pathways with rhamnolipid production in RLQ analysis.** For each rhamnolipid production category (strong-, mild- and non-producers), its correlation value with a single gene or a functional pathway was computed as the dot products between the arrow of phenotypic category and the arrow of the gene or pathway in RLQ axes.

**Supplementary Table 5.** Normalized peak area of metabolomics

**Supplementary Table 6. Read counts of *P. aeruginosa* RNAseq aligned to the PA14 genome**

**Supplementary Table 7. Metadata of RNAseq archived in NCBI SRA database**

## Notes

https://github.com/liaochen1988/Source_code_for_Pseudomonas_Metabolomics_Paper

https://github.com/guisantagui/code_PA_paper

https://github.com/Jinyuan1998/PA_metabolomics_rhamnolipids_SourceCode

### Competing Interest Statement

The authors have declared no competing interest.

### Summary of Updates

We reworked the focus, which was on swarming motility, to the more general problem of the evolution of a secondary metabolic pathway: rhamnolipid biosynthesis. Still the major conclusions stay the same. We conducted new transcriptomics analyses and added Figure 4. We added new results in Figure 5. We also reorganized Figure 1, 2 and 3. Remade Figure 6. We improved the clarity of the manuscript and changed the title to: Evolution and regulation of microbial secondary metabolism.

